# Selective 40S footprinting reveals that scanning ribosomes remain cap-tethered in human cells

**DOI:** 10.1101/806364

**Authors:** Jonathan Bohlen, Kai Fenzl, Günter Kramer, Bernd Bukau, Aurelio A. Teleman

**Affiliations:** German Cancer Research Center (DKFZ), 69120 Heidelberg, Germany; CellNetworks - Cluster of Excellence, Heidelberg University, Germany; Heidelberg University, 69120 Heidelberg, Germany; Heidelberg Biosciences International Graduate School (HBIGS), Germany; National Center for Tumor Diseases (NCT), partner site; Center for Molecular Biology of Heidelberg University (ZMBH) and German Cancer Research Center (DKFZ), DKFZ-ZMBH Alliance, Im Neuenheimer Feld 282, Heidelberg D-69120, Germany

**Keywords:** translation initiation, scanning, ribosome footprinting

## Abstract

Translation regulation occurs largely during initiation. Currently, translation initiation can be studied *in vitro*, but these systems lack features present *in vivo* and on endogenous mRNAs. Here we develop selective 40S footprinting for visualizing initiating 40S ribosomes on endogenous mRNAs *in vivo*. It pinpoints where on an mRNA initiation factors join the ribosome to act, and where they leave. We discover that in human cells most scanning ribosomes remain attached to the 5’ cap. Consequently, only one ribosome scans a 5’UTR at a time, and 5’UTR length affects translation efficiency. We discover that eIF3B, eIF4G1 and eIF4E remain on translating 80S ribosomes with a decay half-length of ∼12 codons. Hence ribosomes retain these initiation factors while translating short upstream Open Reading Frames (uORFs), providing an explanation for how ribosomes can re-initiate translation after uORFs in humans. This method will be of use for studying translation initiation mechanisms *in vivo*.

**HIGHLIGHTS:**

- Selective 40S FPing visualizes regulation of translation initiation on mRNAs *in vivo*
- Scanning ribosomes are cap-tethered in human cells
- Only one ribosome scans a 5’UTR at a time in human cells
- Ribosomes retain eIFs during early translation, allowing reinitiation after uORFs

## INTRODUCTION

Messenger RNA translation is an important step in the gene expression cascade from DNA to RNA to protein, in part because it is the most proximal step to the protein end-product that contributes to a cell’s phenotype. Indeed, mRNA translation efficiency can vary 1000-fold between different mRNAs (Schwanhausser et al., 2011), highlighting the regulatory potential of this process. Much of the regulation of translation happens during translation initiation (Hinnebusch, 2011; Jackson et al., 2010; Pelletier and Sonenberg, 2019; Shirokikh and Preiss, 2018). For instance, activation of 4E-BP blocks recruitment of ribosomes to the 5’ cap (Sonenberg and Gingras, 1998). As another example, many cellular stresses activate the Integrated Stress Response which leads to inactivation of the eIF2-containing ternary complex which normally recruits initiator tRNA to ribosomes (Holcik and Sonenberg, 2005). Furthermore, various 5’UTR sequence elements have been described which affect scanning and initiation, such as upstream Open Reading Frames (uORFs), ribosome shunts, secondary structures such as hairpins and G-quadruplexes, or internal ribosome entry sites (IRESs) (Geballe and Morris, 1994; Leppek et al., 2018; Millevoi et al., 2012; Mitchell and Parker, 2015).

One category of regulatory elements in 5’UTRs are uORFs. uORFs can be classified into those with weak Kozak sequences and those with strong ones. The ones with weak Kozak sequences do not strongly impact ribosomes as they scan for the main ORF on the transcript, because they simply scan past the uORF in a process termed leaky scanning. The ones with strong Kozaks, however, pose a challenge because ribosomes recognize them as bona fide translation start sites and therefore translate the uORF. Upon terminating translation of the uORF, they need to re-initiate translation on the main ORF in order to express the encoded protein. Translation re-initiation is a process that is currently not well understood, but probably involves stabilization of the ribosome on the mRNA, re-recruitment of an initiator tRNA, and resumed scanning of a ribosomal complex that is competent to initiate another round of translation. How this happens mechanistically is not known, yet it happens frequently. Nearly half of all human mRNAs contain at least one uORF (McGillivray et al., 2018) and ribosome footprinting experiments have revealed that many of these are translated *in vivo* (Ingolia et al., 2011). Hence there is a need to better understand the process of translation re-initiation. One aspect worth pointing out is that translation re-initiation appears to be different in human cells versus yeast cells (Kozak, 2001). In humans, most uORFs are permissive for reinitiation, meaning they allow translation of the main ORF downstream to occur, whereas in yeast most uORFs are not (Jackson et al., 2012; Johansen et al., 1984; Kozak, 1984; Liu et al., 1984; Yun et al., 1996). Instead, the best studied example of translation re-initiation in yeast is on the GCN4 mRNA, and this requires specific cis-acting elements flanking the uORFs to stabilize the ribosome on the mRNA and to permit reinitiation (Mohammad et al., 2017; Szamecz et al., 2008). Hence some molecular details of translation initiation which permit reinitiation in human cells may be different from yeast.

Generally, translation can be studied using two major types of approaches. The first are *in vitro* systems consisting either of cell-free translation extracts or reconstituted systems. These systems have incredible power and resolution at deciphering individual steps of translation as well as the molecular functions of individual proteins such as initiation factors. Indeed, much of our understanding of the succession of steps that constitutes translation derives from such systems (Hinnebusch, 2011; Jackson et al., 2010, 2012). These systems, however, have the disadvantage that they usually analyze translation on short synthetic mRNA rather than endogenous mRNAs, and hence lack many of the interesting and complex features, both known and unknown, of endogenous mRNAs. Secondly, since these systems are cell-free, they also lack components and regulatory mechanisms that are present in a cell. The second major approach, based on RNA protection assays by Joan Steitz’ lab (Steitz, 1969), is ribosome footprinting (Ingolia et al., 2009; Ingolia et al., 2011), which allows the localization and quantification of translating 80S ribosomes *in vivo* in a cell on endogenous mRNAs. This approach, however, cannot resolve 40S ribosomes because they are more loosely associated to the mRNA than translating 80S ribosomes, hence it cannot visualize the regulatory steps occurring during translation initiation. Furthermore, it cannot identify when and where translation factors are acting on the ribosome. To study translation regulation, it would be useful to visualize initiation of translation in the cell on endogenous mRNAs at single nucleotide resolution on single transcripts. Furthermore, if it were possible to detect when individual initiation factors join the ribosome and then disengage during the initiation process, this would indicate when these initiation factors are acting. Together, this would provide a detailed understanding of the orchestrated succession of steps that constitute initiation on an mRNA of interest.

We develop here selective 40S footprinting to visualize the successive steps of translation initiation *in vivo* on endogenous mRNAs, and to pinpoint when translation initiation factors join the ribosome to act, and then later disengage. (Please note that for simplicity we use here the term “40S” to denote all variants of the 40S, such as 43S and 48S ribosomes). Using this technology, we find that in human cells 5’UTR scanning happens in a cap-tethered fashion whereby the scanning 40S ribosome remains attached to eIF3B, eIF4G1, eIF4E and the mRNA cap up to the start codon of the main ORF. This implies that only one ribosome can scan a 5’UTR at one time in human cells, making 5’UTR length a limiting factor for optimal translation efficiency. Furthermore, we find that a substantial portion of initiation factors perdure on early 80S translating ribosomes, with a half-length of ∼12 codons. Consistent with this, we see that ribosomes retain eIF3B, eIF4G1 and eIF4E when they reach the stop codon of translated uORFs, which are usually much shorter than 36 nt. This likely explains their ability to remain attached to the mRNA and resume scanning after uORF translation termination. Finally, we see that eIF2 disengages from 40S ribosomes at start codons of main ORFs and uORFs, and is quickly re-recruited to the ribosome after termination on a uORF. Together, this provides a molecular sequence of events which explains how translation reinitiation can occur after a short uORF, but not after main ORFs which are significantly longer. This technology we have developed will be of use for dissecting mechanisms of translation initiation *in vivo* in the future.

## RESULTS

### 40S footprinting visualizes scanning ribosomes in vivo

To study translation initiation *in vivo*, we aimed to visualize ribosomes scanning on cellular mRNAs, and to identify at which point during this process individual initiation factors join the ribosome to act, and then leave. Since 40S ribosomes are weakly associated with the mRNA, they are not retained in standard ribosome footprinting approaches. Hence, we first adapted for human cells “translation complex profile sequencing” (TCP-seq), a version of ribosome footprinting that includes a crosslinking step to stabilize scanning 40S ribosomes on the mRNA (Archer et al., 2016; Shirokikh et al., 2017). We tested in HeLa cells different concentrations and combinations of para-formaldehyde (PFA) and dithiobis(succinimidyl propionate) (DSP) for crosslinking. We added the DSP to the TCP-seq protocol to further stabilize protein-protein interactions. This revealed that insufficient crosslinking failed to stabilize 40S-mRNA complexes, whereas excessive crosslinking caused ribosomes to aggregate, leading us to find an optimal concentration of 0.025% PFA + 0.5 mM DSP which does not produce large ribosome aggregates (as observed on a polysome gradient, Suppl. Fig. 1A) yet stabilizes both ribosomes and initiation factors on mRNA (Suppl. Fig. 1B) without cross-linking non-specific interactions (Tubulin, Suppl. Fig. 1B). After *in* vivo crosslinking, cell lysis and RNAse digestion, we separated scanning 40S ribosomes from elongating 80S ribosomes on a sucrose gradient and sequenced the footprints of these two populations (Fig. 1A). As expected, 40S footprints localize predominantly to 5’UTRs whereas 80S footprints are mainly found in open reading frames (ORFs) (Fig. 1B). This allows for an unprecedented mRNA-resolved transcriptome-wide view of scanning ribosomes in human cells. For instance, for the eIF5A mRNA, scanning 40S ribosomes are detected in the 5’UTR which are then converted to elongating 80S ribosomes at the ORF start (Fig. 1C). After translation termination on main ORFs, 40S ribosomes could either fall off, or scan off the 3’end of the mRNA (Bertram et al., 2001; Dever and Green, 2012). Since we see essentially no 40S ribosomes in 3’UTRs (Fig. 1B-C), this means they are falling off the mRNA at the stop codon. As observed in yeast (Archer et al., 2016), 40S footprints have a size distribution distinct from that of elongating 80S footprints (Fig. 1D-E). A 2-dimensional metagene plot, which resolves footprint position on the x-axis and footprint length on the y-axis, of 40S footprints relative to all main-ORF start codons (“start codon metagene plot”) shows that scanning ribosomes (positions −100 to −40 relative to the start codon) have heterogeneous footprint sizes ranging from 20nt to 80nt (Fig. 1D). Also as expected, 40S footprints show no triplet periodicity (Fig. 1D), whereas 80S footprints do (Fig. 1E). When scanning ribosomes reach the start codon, they pause, as observed by a strong enrichment of 40S footprints overlapping the start codon, showing that initiation is significantly slower than scanning *in vivo* in human cells. Two main populations of footprints can be observed overlapping the start codon, with footprint lengths of circa 20 and 30 nt. Each of these has a ‘tail’ that extends diagonally up and to the left, representing a series of footprints that have the same 3’ end but varying 5’ ends (bottom of Fig. 1D). This suggests that the 5’ ends of initiating 40S ribosomes protect mRNA less robustly than the 3’ ends. In sum, our modified TCP-seq enables the investigation of ribosome scanning and initiation in human cells.

**Figure 1:**
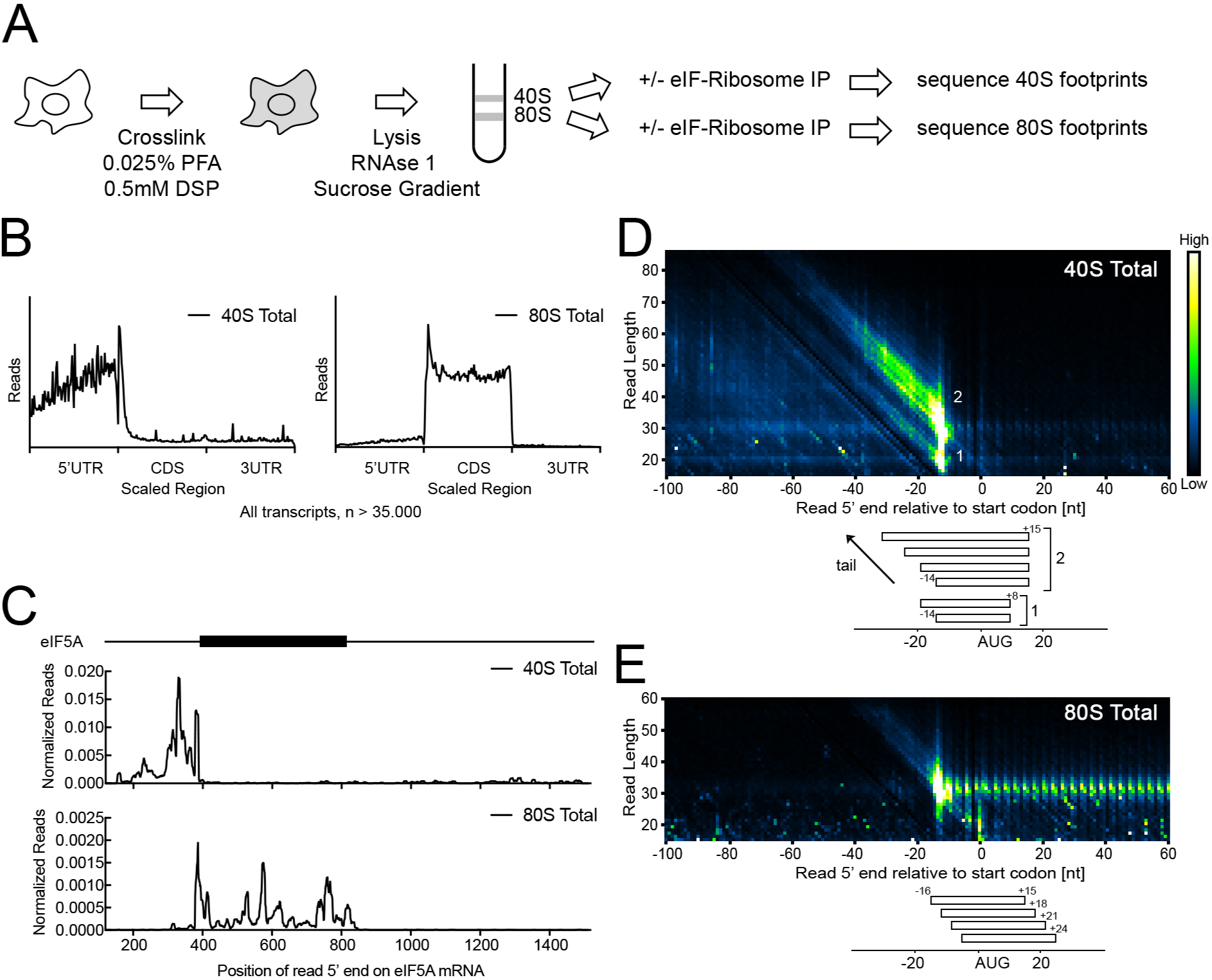
40S ribosome footprinting in human cells. **(A)** Schematic diagram illustrating selective and total 40S and 80S ribosome footprinting in human cells. HeLa cells are crosslinked, lysed and RNAse treated. 40S and 80S fractions are separated on a sucrose gradient and then immunoprecipitation of eIF-bound ribosomes is carried out. RNA is extracted and footprints are deep-sequenced. **(B)** 40S ribosome footprints are located in 5’UTRs but not ORFs or 3’UTRs. Metagene plot of all 40S (left) or 80S (right) reads mapped to all human protein coding transcripts with 5’UTR length > 33 nucleotides (n = 35.921). **(C)** 40S and 80S ribosome footprint distribution on eIF5A mRNA (ENST00000336458). Curves smoothened with sliding window of 10 nt. Black box = eIF5A coding sequence. **(D-E)** 40S ribosome footprinting reveals stalling and processing of 40S ribosomes on translation start sites. Top panels: Metagene plot of length resolved 40S (D) or 80S (E) ribosome footprints aligned to the main ORF start codon on >41.000 human transcripts. The number of reads is displayed according to a linear color scale, shown on the right. Bottom panels: Schematic representation of footprint species of different lengths on the start codon.

### Selective 40S footprinting reveals steps of translation initiation in vivo

We next combined this with an immunoprecipitation step to isolate ribosomes that contain particular proteins of interest, similar to selective ribosome profiling on 80S ribosomes (Oh et al., 2011; Schibich et al., 2016). We immunoprecipitated 40S ribosomes that contain endogenous eIF2S1, eIF3B or eIF4G1 (Suppl. Fig. 2A-C). EIF2S1/eIF2*α* is a component of the ternary complex, which is responsible for recruiting initiator tRNA to initiating ribosomes. eIF3B is a core subunit of the eIF3 complex, which promotes attachment of scanning ribosomes to mRNA and serves as a central docking platform for many initiation factors. eIF4G1 is the core scaffold of the eIF4F complex which links eIF4E and hence the 5’cap to the ribosome (Jackson et al., 2010). This successfully identified different subpopulations of 40S ribosomes: a metagene plot relative to main ORF start codons showed that ribosomes overlapping the start codon are partially depleted of eIF2 (Suppl. Fig. 2D, which is normalized to library size, and Fig. 2A which normalizes the selective footprints down to the total footprints in the scanning region). This is expected because the eIF2*α*-containing ternary complex is evicted prior to 60S subunit joining (Kapp and Lorsch, 2004; Pisarev et al., 2006; Unbehaun et al., 2004), hence 40S ribosomes spend part of their time on the start codon with eIF2, and part of their time without eIF2. This selective footprinting allows annotation of the various populations that can be observed in the 2-dimensional metagene plot (Fig. 2B). Scanning ribosomes (population 1) contain eIF2*α* (Fig. 2C-C’), whereas 40S ribosomes overlapping the start codon (2 and 3) are partially depleted of eIF2*α*. Scanning ribosomes containing eIF3B (Fig. 2D) have larger footprints than the average scanning 40S ribosome (the upper region is green and the lower region is red in the ratiometric image Fig. 2D’), indicating that part of the RNA protection of scanning ribosomes is caused by the large 600–800 kDa eIF3 complex (Erzberger et al., 2014). Analysis of ribosomes on the start codon (populations 2 & 3) revealed that eIF3B is enriched in the large ‘tail’ of populations 2 & 3 (Fig. 2D’), representing footprints with extended RNA protection on the 5’ end but unchanged 3’ends. This indicates that the eIF3 complex protects loosely a region of up to 40nt on the 5’ end of the 40S ribosome. Since no structure is available for eIF4F on the ribosome, it has been unclear on which side of the ribosome this complex sits (Shirokikh and Preiss, 2018). We find that the presence of eIF4G1 causes protection on the 5’ end of the ribosome (Fig. 2E-E’). These data suggest eIF4F contacts the mRNA when it exits the ribosome, and hence the helicase may function by pulling mRNA through the ribosome rather than pushing it in through from the front. The 20-nt footprints in population 3 may represent an intermediate step of translation initiation, as has been observed in yeast (Archer et al., 2016), or a conformation where the A-site is not occupied and hence accessible to RNAse (Wu et al., 2019). In sum, selective 40S footprinting enables us to localize scanning 40S ribosomes on individual mRNAs and to identify when initiation factors join or disengage from the scanning 40S.

**Figure 2:**
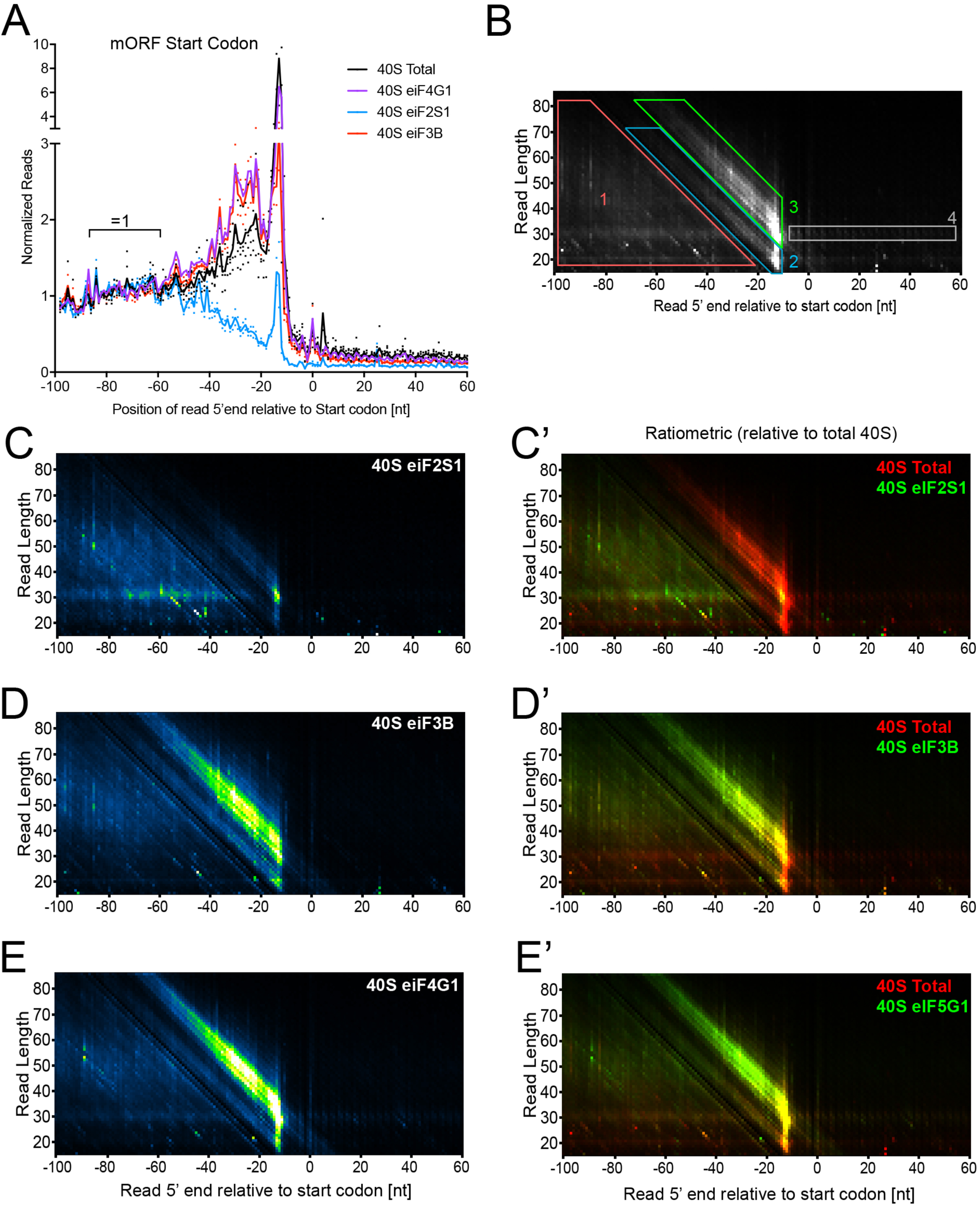
Selective 40S ribosome footprinting in human cells. **(A)** eIF2S1 dissociates from the ribosome during start codon recognition. Metagene plot of total 40S and eIF3B-, eIF4G1-, or eIF2S1-selective 40S ribosome footprints aligned to the start codon of all human protein coding transcripts (n = 41.244). Reads are mapped to the position of the read 5’ end. Curves are scaled to total read counts in the region −98 to −69 in front of the start codon. Curves show the average of 2-4 biological replicates (dots). Graphs normalized to library size are shown in Suppl. Fig. 2D. **(B)** Illustration of distinct 40S ribosome populations visible in the 2D start codon metagene plot. (1) scanning ribosomes, (2) and (3) 40S ribosomes on start codon, (4) background signal from 80S ribosome disassembly during sample preparation. **(C-E’)** Length resolved start codon metagene plots of 40S footprints from eIF2S1 (C-C’), eIF3B (D-D’), or eIF4G1 (E-E’) containing 40S ribosomes. Intensity scales are adjusted to the normalization in (A). Panels (C’, D’ E’) show ratiometric images of selective footprints (green) versus total 40S footprints (red).

### Scanning of the 5’UTR mainly occurs in a cap-tethered fashion

The 5’ cap of mRNAs recruits ribosomes onto the mRNA via a bridge of interactions from the cap to eIF4E, to eIF4G1, to eIF3, to the 40S ribosome. As ribosomes scan from the cap towards the main ORF start codon, they could either let go of the 5’cap by severing one of these interactions, or they could remain attached to it (Fig. 3A). In the latter case, the 5’UTR would have to loop. Which of these two options happens *in vivo* is not known, but this issue has important conceptual and functional consequences for translation regulation (Jackson et al., 2010; Shirokikh and Preiss, 2018). We reasoned that eIF4E-selective 40S footprinting should allow us to address this longstanding open issue. If ribosomes are severed from eIF4E during scanning, they should become depleted of eIF4E as they scan in the 3’ direction, causing a drop in density of eIF4E-selective footprints (Fig. 3A). If instead they remain tethered to eIF4E, eIF4E-selective footprint densities should remain uniform throughout the 5’UTR. To distinguish these two possibilities, we performed eIF4E-selective 40S footprinting (Suppl. Fig. 3A-B). Surprisingly, a metagene plot of 5’UTRs, where each 5’UTR length is scaled to 100%, shows that ribosomes retain eIF3B, eIF4G1 and eIF4E in constant proportion throughout the entire scanning process from the cap to the main ORF start codon (Fig. 3B). In agreement with this, a metagene profile of all main start codons shows that 40S ribosomes stoichiometrically retain eIF4G1 and eIF4E up to the start codon of the main ORF (Fig. 3C and Suppl. Fig. 3D). Together, these data indicate that in most cases ribosomes remain associated to eIF4E during scanning up to the start codon in human cells *in vivo*. It may be that a small percentage of ribosomes let go of eIF4E at the start codon itself (14% drop in area under the curve from −45 to −5 for the eIF4E-selective profile compared to total 40S, Fig. 3C), however this does not fit with the quantitative retention of eIF4E throughout 5’UTRs (Fig. 3B) and the fact that 50% of all 5’UTRs have uORFs in them: If eIF4E were lost at start codons, this would happen on translated uORFs thereby causing a reduction in eIF4E binding in the 5’UTR.

**Figure 3:**
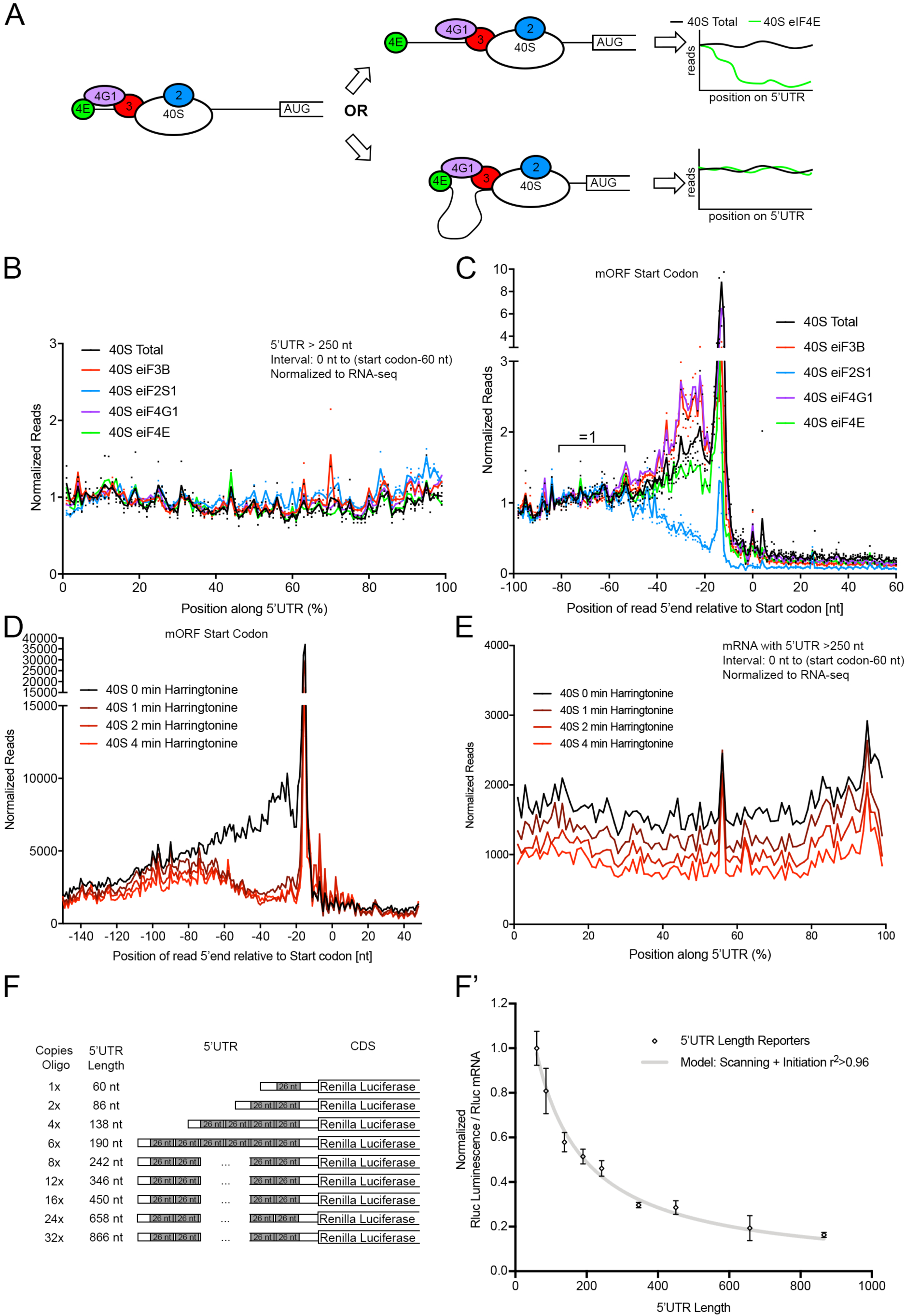
Only one cap-tethered 40S ribosome scans a human mRNA 5’UTR at one time. **(A)** Schematic diagram of cap-severed versus cap-tethered scanning and how they can be differentiated by eIF4E selective 40S ribosome footprinting. **(B)** Scanning 40S ribosomes are bound to eIF2S1, eIF3B, eIF4G1 and eIF4E in equal proportions throughout the entire 5’UTR. Metagene plot of footprints on all 5’UTRs longer than 250nt (n = 13.439). Position along 5’UTR is scaled from 0% (5’ cap) to 100% (start codon – 60nt). The last 60 nucleotides are excluded because they contain initiating ribosomes (shown in panel C). The five curves were normalized to each other using the scaling factors from Figure 2A, and to RNA-seq read counts at each position to account for varying mRNA abundance along the 5’UTR due to alternative transcription start sites. Curves show average of 2-3 biological replicates (dots). **(C)** Initiation factors eIF3B, eIF4G1 and eIF4E are retained on 40S ribosomes on main ORF start codons. Start codon metagene plot of total and selective 40S ribosome footprints on all human protein coding transcripts (n = 41.244). Reads are mapped to the position of the read 5’ end. Curves are normalized to each other using the scaling factors from Figure 2A. Curves show average of 2-3 biological replicates (dots). **(D-E)** Harringtonine block of initiating 80S ribosomes does not cause 40S queuing in front of the start codon (D), but instead causes depletion of 40S ribosomes in 5’UTRs (E). (D) Start codon metagene plot of total 40S ribosome footprints on all human protein coding transcripts (n = 41.244) at different timepoints after harringtonine treatment (2 µg/ml, 37°C). Reads are mapped to the position of the read 5’ end. Counts are normalized to the number of mapped reads in the library. (E) Metagene plot of total 40S ribosome footprints along 5’UTRs with length > 250nt (n = 13.439). Position along 5’UTR scaled from 0%-100% as in panel B. Read counts were normalized to the number of mapped reads in each library and to RNA-seq read counts at each position to account for varying mRNA abundance along the 5’UTR due to alternative transcription start sites. **(F-F’)** 5’UTR length limits the translational output of mRNAs in human cells. Renilla luciferase luminescence normalized to mRNA levels (F’) for translation reporters containing 5’UTRs of varying lengths illustrated in (F). 5’UTRs were created by inserting multiple copies of an unstructured 26-mer in front of the renilla luciferase ORF. Renilla luciferase mRNA levels were normalized to Actin B mRNA levels. One representative example of three biological replicates is shown.

It is possible that the interaction of eIF4E with the mRNA cap is severed during scanning, although unlikely since binding of eIF4E to eIF4G1 increases eIF4Es affinity for the cap (von Der Haar et al., 2000). To test this, we treated cells with harringtonine and performed 40S footprinting. Harringtonine specifically arrests initiating 80S ribosomes on the start codon (Fresno et al., 1977; Ingolia et al., 2012). If ribosomes release the cap during scanning, the cap should be free to recruit multiple new rounds of 40S ribosomes, which should accumulate on the 5’UTR, likely amassing in front of the stalled 80 ribosome. Hence 40S footprints should increase with harringtonine treatment over time, and we would expect 40S peaks forming in front of the start codon. If instead the ribosome on the start codon remains cap-tethered, this will block recruitment of another ribosome to the mRNA. Hence over time, 5’UTRs should become progressively depleted of 40S ribosomes, and the number of footprints in the 5’UTR should diminish. Indeed, the latter was the case. Treatment with harringtonine, which led to an arrest of initiating 80S ribosomes and a run-off of elongating 80S ribosomes (Suppl. Fig. 3 E-F), did not cause an accumulation of 40S ribosomes upstream of the main start codon (Fig. 3D). In fact, it caused a reduction. Furthermore, there was a progressive depletion overall of 40S footprint reads on 5’UTRs (Fig. 3E). Also consistent with this is the fact that harringtonine treatment causes ribosomes to accumulate into a monosome peak on a sucrose gradient, indicating the presence of a single 80S ribosome stalled on the start codon, rather than heavier complexes corresponding to one 80S ribosome plus multiple 40S ribosomes on the 5’UTR. Although such polysome gradients are usually run in the absence of crosslinkers, which would cause a loss of 40S-containing complexes on the mRNAs, we also see this with our cross-linking conditions which stabilize scanning 40S ribosomes (Suppl. Fig. 3G). From these data we conclude that indeed most 40S ribosomes do not let go of the mRNA cap during scanning, i.e. scanning is mainly cap-tethered in human cells.

A consequence of cap-tethered scanning is that 5’UTR length should affect mRNA translation efficiency. The time it takes to scan an entire 5’UTR puts a minimum limit on how quickly a new ribosome can be recruited to the mRNA, and hence how quickly a new round of protein synthesis can be initiated. This implies that all things equal, the longer the 5’UTR of an mRNA, the lower its translation efficiency should be. To test this, we synthesized a series of translation reporters containing renilla luciferase (RLuc) with 5’UTRs of homogeneous quality but increasing length, generated by multimerizing a 26-mer sequence that lacks secondary structure or uORFs (Fig. 3F). We then transfected HeLa cells with these constructs and quantified RLuc activity normalized to the amount of reporter mRNA in the cells. This revealed that indeed the longer the 5’UTR, the lower the RLuc expression from these mRNAs (Fig. 3F’ and Suppl. Fig. 3H). In fact, the data fit very well (r>0.96) to a simple model whereby the amount of time it takes to scan an mRNA (t) is equal to the length of the 5’UTR (l) divided by a scanning velocity (v), plus a fixed time (c) for initiation on the ATG (t=l/v + c) and hence the number of initiations per unit time is 1/t (Fig. 3F’). Altogether, these data validate the finding that in human cells ribosomes scan mainly in a cap-tethered manner, which has functional consequences, making 5’UTR length one important parameter in determining translation efficiency of an mRNA.

### Initiation factors eIF3B, eIF4G1 and eIF4E persist on elongating 80S ribosomes with a decay half-length of ∼12 codons

Since we observed that eIF3B, eIF4G1 and eIF4E are retained on 40S ribosomes up to the start codon, we asked if they can remain bound to 80S ribosomes after subunit joining. To this end, we performed selective 80S footprinting, immunoprecipitating each of these initiation factors from 80S fractions of a sucrose gradient (Suppl. Figs. 2A-C, 3A). Interestingly, sequencing of these footprints revealed a strong enrichment of eIF3B-, eIF4G1- and eIF4E-containing 80S ribosomes on main ORF start codons, compared to total 80S ribosomes (Suppl. Fig. 4A). In contrast, eIF2S1 is de-enriched on 80S ribosomes (Suppl. Fig. 4A), consistent with it falling off at the start codon (Fig. 2A). To analyze the dissociation of these initiation factors from 80S ribosomes as they elongate on the ORF, we normalized down the height of the selective 80S footprint peaks to be equal to the total 80S on the start codon (Fig. 4A). Interestingly, this revealed that initiation factors do not dissociate immediately from 80S ribosomes as they start elongating, but rather dissociate over time (Fig. 4A). By calculating the ratio of selective versus total 80S footprints for each position on the ORF, we found an exponential dissociation of eIF3B, eIF4G1 and eIF4E from the 80S ribosome, with a half-length of circa 36 nucleotides (Fig. 4B) corresponding to roughly 12 cycles of elongation. The good fit to an exponential decay curve (r^2^>0.82) suggests this dissociation process is stochastic in nature. We therefore conclude that a significant proportion of eIF3 and eIF4 complexes remain associated to initiating and early translating 80S ribosomes. This has several functional consequences which we analyze below.

**Figure 4:**
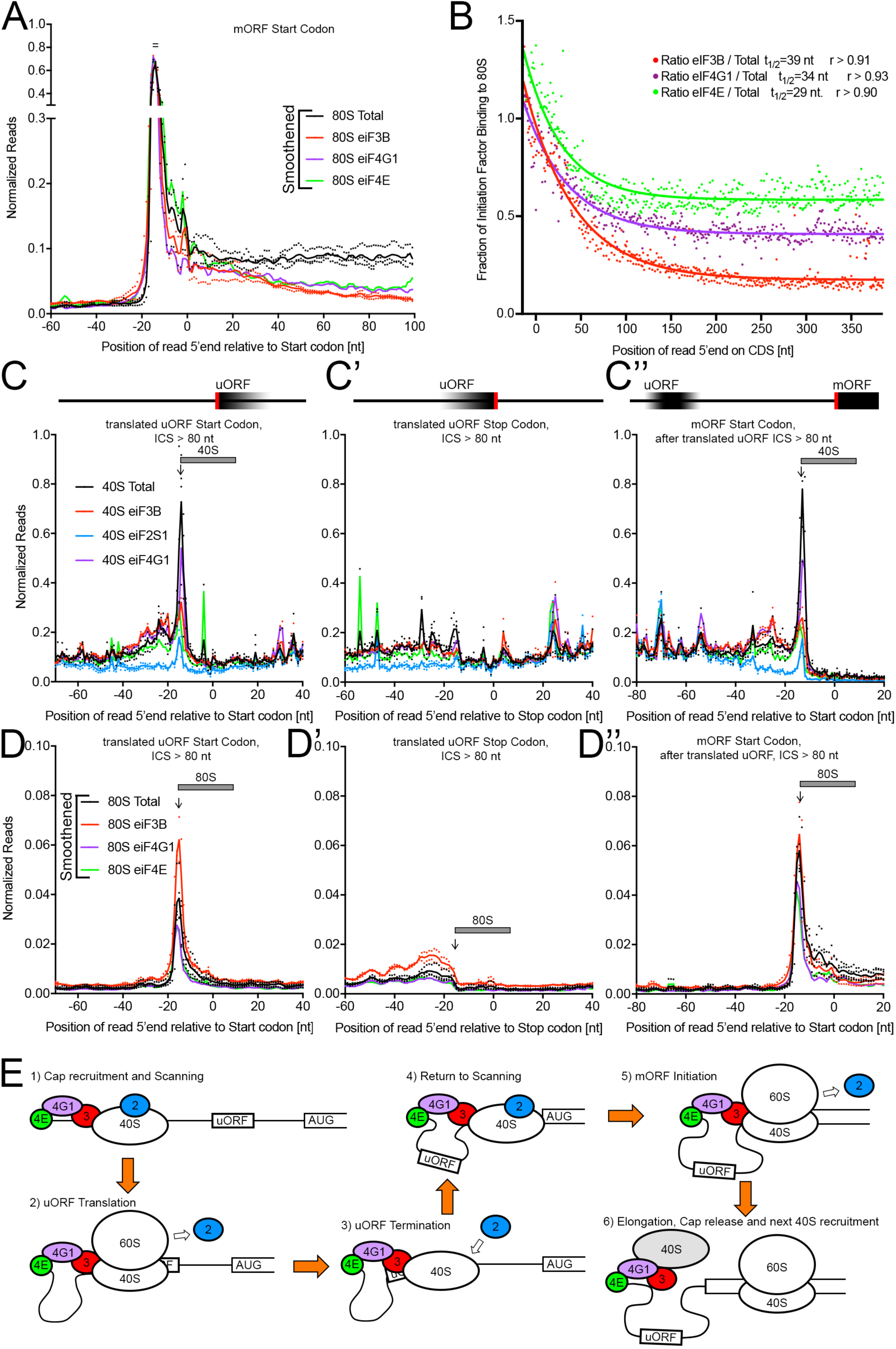
Persistent binding of eIF3B, eIF4G1 and eIF4E on early elongating 80S ribosomes leads to initiation factor retention after translation of uORFs. **(A)** eIF3B, eIF4G1 and eIF4E selective 80S ribosome footprints are enriched on start codons and then progressively de-enriched. Start codon metagene plot of total and selective 80S ribosome footprints on all human protein coding transcripts (n = 41.244). Reads are mapped to the position of the read 5’ end. Counts are normalized to the height of the peak on the start codon. Graphs normalized only to library size are shown in Suppl. Fig. 4A. **(B)** Initiation factor release from translating 80S ribosomes follows an exponential decay. Ratio of selective 80S ribosome footprints to total 80S ribosome footprints along the coding sequences of all protein coding mRNAs (n = 41.244). Reads are mapped to 5’ end and offset by +15 nucleotides. Read counts are normalized as in (A). The exponential decay function 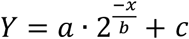, where b is the decay half-length, was fit to the data points.

**(C-D’’)** Initiation factor binding to ribosomes persists through the translation of uORFs. Metagene plots of total and selective 40S footprints (C-C’’) or 80S footprints (D-D’’) relative to the start codon (C and D) or stop codon (C’ and D’) of translated uORFs (n = 6663), or to the start codon of the main ORF downstream (C’’ and D’’). Reads are mapped to the position of the read 5’ end. Curves are normalized to each other using the scaling factors from Figure 2A (for 40S) or Figure 4A (for 80S).

**(E)** Model summarizing our data of translation reinitiation.

Curves in panels A, C-C’’ and D-D’’ show the average of 2-3 biological replicates (dots).

### Initiation factors persist past uORFs on translating ribosomes

Roughly half of all human mRNAs contain upstream Open Reading Frames (uORFs) and ribosome footprinting experiments have shown that many of these are translated (Calvo et al., 2009; Ingolia et al., 2011; Johnstone et al., 2016). In such cases, ribosomes need to terminate translation on the uORF, resume scanning, and re-initiate downstream on the main ORF, a process called translation re-initiation (Gunisova et al., 2018; Jackson et al., 2012). The molecular steps of translation re-initiation are not fully understood. In particular, it is unclear how ribosomes re-recruit initiation factors after uORF translation. Our eIF selective footprinting data provide a possible molecular explanation for this: if the uORFs are short enough, initiation factors such as eIF3 and eIF4G might be retained on the 80S ribosome up to the uORF stop codon. This would make it competent to reinitiate by stabilizing association with the mRNA and providing a platform to re-recruit the initiation factors that are lost at 60S joining such as eIF1, 1A, 2 and 5. To study this, we analyzed uORFs with detectable 80S footprints on their start codons, indicating that they are translated. From these, we selected only uORFs with an intercistronic spacing (ICS) ≥80 nucleotides to the main ORF start codon, in order to resolve uORF translation from main ORF translation (6663 uORFs in 3916 transcripts). Metagene plots of such uORFs confirmed that both 40S ribosomes and 80S ribosomes accumulate on their start codons (Fig. 4C and 4D), indicating they are recognized as translation start sites and then translated. Indeed, 80S ribosomes translating these uORFs display triplet periodicity when the curves are not smoothened (Suppl. Fig. 4B-C). Just like on main-ORF start codons, 40S ribosomes on these “translated uORF” start codons have eIF3B, eIF4G1 and eIF4E, but are depleted of eIF2S1 (blue trace is below the others, Fig. 4C). Note that all 40S and 80S graphs in Fig. 4 are normalized using the same values as the ones used in Fig. 2A and Fig. 4A, respectively, to make them directly comparable. Also worth pointing out is that the coding sequences of the 6663 uORFs in this metagene plot have varying lengths (ranging from 6nt to 1284nt, with a mean of 36nt), hence although the start codons are aligned, the stop codons are not aligned, and occur at various downstream positions. As a result, scanning 40S ribosomes and eIF2S1 are depleted directly after the start codon, but they progressively return to baseline further downstream as the uORFs asynchronously finish (Fig. 4C). If we align translated uORF stop codons, several observations can be made (Fig. 4C’ and 4D’). Firstly, we see that 80S ribosomes disappear and scanning 40S ribosomes re-appear downstream of the stop codon, suggesting that reinitiation is performed by scanning 40S ribosomes, not 80S ribosomes. Re-initiation was originally proposed to happen via scanning 40S ribosomes, but to our knowledge this was never directly proven, and recent work suggested that scanning 80S ribosomes might be responsible for re-initiation (Zhou et al., 2018). Secondly, one sees that in the uORF coding sequence eIF2S1 is depleted, but is quickly re-recruited to 40S ribosomes after termination on the uORF stop codon (blue trace re-joins the other traces at the stop codon, Fig. 4C’). Thirdly, one sees that after the uORF stop codon the other initiation factors eIF3B, eIF4G and eIF4E are not depleted from 40S ribosomes. Instead, relative to total 40S ribosomes, these initiation factors are present at similar levels after the stop codon as they were before the uORF start codon (Fig. 4C-C’). Indeed, as these 40S ribosomes move downstream and reach the main ORF start codon (Fig. 4C’’ and 4D’’), they have eIF3B, eIF4G1 and eIF4E, they become depleted of eIF2S1, and they convert to 80S ribosomes, just as on transcripts lacking uORFs. Since most translated uORFs have short coding sequences, these data are consistent with a model whereby most initiation factors are retained on 80S ribosomes as they translate uORFs, thereby enabling the post-termination 40S ribosomes to re-recruit eIF2 and to recommence scanning (Fig. 4E). In contrast, on main ORFs which tend to be significantly longer than 36nt, the initiation factors are likely depleted by the time the elongating 80S reaches the stop codon, causing ribosomes to release the mRNA after terminating and to not scan the 3’UTR (Fig. 1B).

### Cellular stress induces low tRNA-iMet binding to the ternary complex, not low ternary complex binding to 40S

Inactivation of eIF2 is one of the principal modes of translation regulation. Cellular stresses such as oxidative stress, proteotoxic stress, dsRNA, or low amino acids, lead to phosphorylation of eIF2S1/eIF2*α* and consequently inactivation of the eIF2 complex (McConkey, 2017; Pakos-Zebrucka et al., 2016). On the one hand, this causes a global drop in translation. On the other hand, it activates expression of stress response proteins such as ATF4 via a re-initiation mechanism. Human ATF4 has three uORFs in its 5’UTR. In brief, the canonical model of ATF4 regulation is that ribosomes translate the uORFs and then recharge with the eIF2•GTP•inititator-tRNA ternary complex. If eIF2 activity is high, they recharge quickly and translate a decoy uORF. If they recharge slowly, they skip past the decoy uORF and translate the ATF4 main ORF (Harding et al., 2000; Lu et al., 2004; Vattem and Wek, 2004). Hence the speed of recruitment of the ternary complex is key in this regulatory mechanism, which may also apply to other genes with overlapping uORFs. *In vitro*, it has been shown that two different modes of initiator tRNA recruitment are possible (Sokabe et al., 2012). Initiator tRNA can first bind eIF2•GTP to form the ternary complex. This ternary complex then binds the ribosome. Alternatively, eIF2 can first bind the ribosome, and then recruit initiator tRNA. Which of these two happens predominantly *in vivo* is not known. To test this, we performed eIF2S1-selective 40S footprinting on cells treated with tunicamycin to induce ER stress (Suppl. Fig. 2B). We then analyzed eIF2 recruitment after translated uORFs. As expected, both in the presence and absence of stress, eIF2*α* is depleted from 40S ribosomes on uORF start codons (blue and pink curves, Fig. 5A). Interestingly, eIF2*α* is equally quickly re-recruited after translated uORF stop codons both in the unstressed and stressed conditions (Fig. 5A’). Instead, the amount of initiator-tRNA detected in our eIF2S1-selective 40S pull-downs was reduced in the presence of stress (Fig. 5B). Additionally, 40S ribosomes initiating on the main ORF downstream of the uORF are mildly reduced in the stress condition (Fig 5A’’). Together, these data indicate that in the presence of stress, eIF2 is re-recruited to re-initiating 40S ribosomes just as quickly as in the absence of stress, however these eIF2 complexes contain less initiator tRNA. The eIF2•40S complex then recruits initiator tRNA, and this happens in a delayed manner in the presence of stress. This thereby provides an *in vivo* view of how eIF2 and the initiator tRNA are recruited to re-initiating 40S ribosomes.

**Figure 5:**
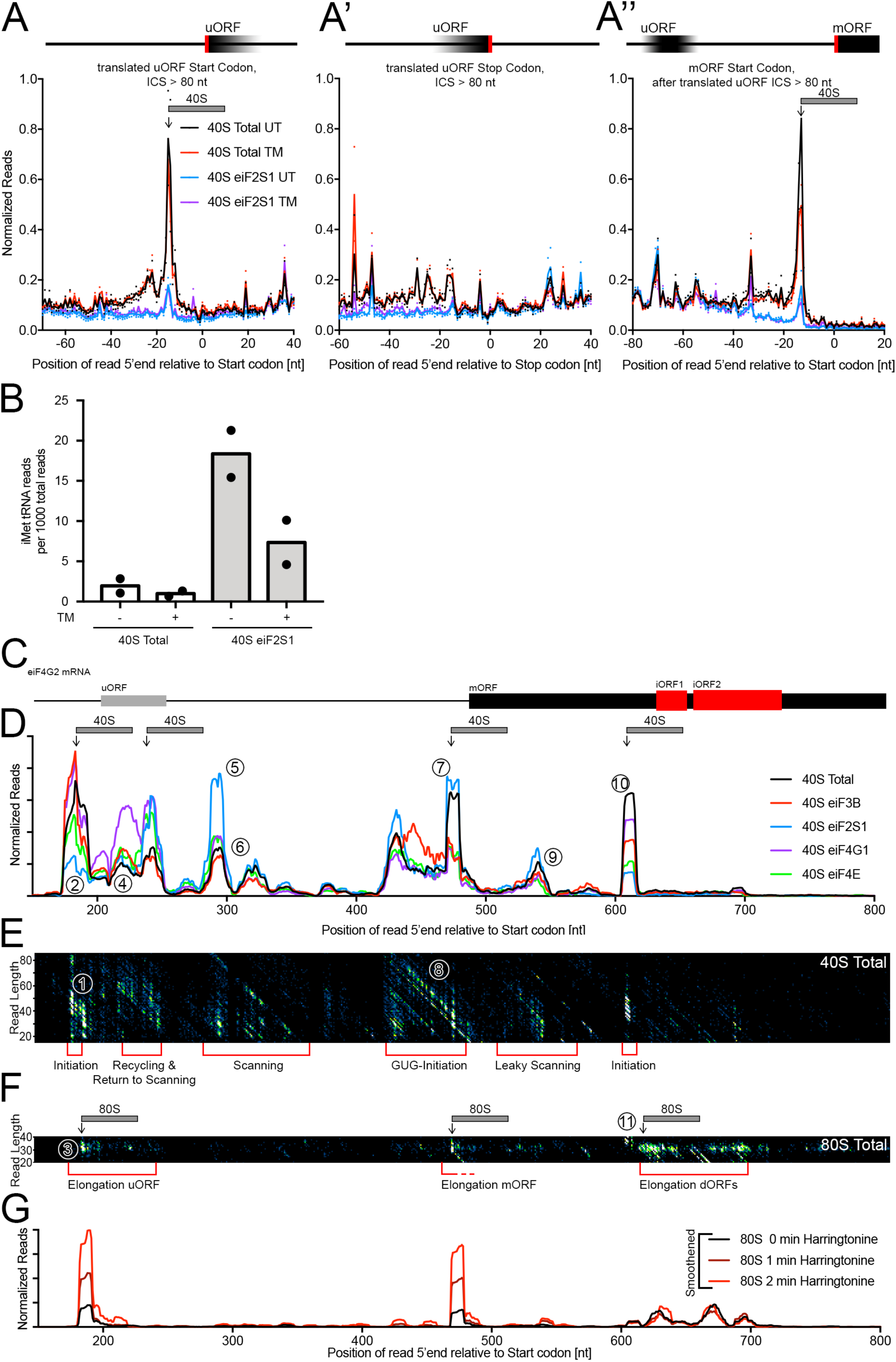
Initiation factor 2 complex is slowly recharged with iMet-tRNA on 40S ribosomes during stress and can initiate translation on near-cognate start codons. **(A-A’’)** Initiation factor eIF2S1 is re-recruited to scanning 40S ribosomes after translation of a uORF independent of cellular stress. Metagene plots of total or selective 40S footprints relative to the start codon (A) or stop codon (A’) of translated uORFs (n = 6663), or to the start codon of the main ORF downstream (A’’). Cells were either untreated (“UT”, black and blue curves) or treated with tunicamycin (replicate 1: 250 ng/ml, replicate 2: 1 µg/ml) for 16 hours. Reads are mapped to the position of the read 5’ end. Graphs are normalized to number of scanning 40S ribosome footprints at positions −98 to −69 in front of the start codon of an all transcript metagene profile (analogous to normalization in figure 2A). **(B)** Binding of methionine initiator-tRNA to eIF2S1 associated ribosomes is stress dependent. Count of iMet tRNA reads per 1000 sequenced reads in total 40S footprint and eIF2S1-selective 40S footprint libraries from untreated and tunicamycin treated cells. **(C)** Diagram illustrating positions of ORFs in the eIF4G2 mRNA 5’ region (ENST00000339995). **(D-E)** Reinitiation, near-cognate GTG-initiation and leaky scanning occur on the eIF4G2 mRNA. (D) Counts of total and selective 40S ribosome footprints along the 5’ region of the eIF4G2 mRNA. Reads are normalized using the scaling factors from Fig. 2A. Curves are smoothened with a 10 nt sliding window. (E) Same data as in C, but length-resolved. **(F)** Length resolved footprint distribution of total 80S footprints on eIF4G2 mRNA proximal region. **(G)** Harringtonine treatment leads to 80S accumulation at uORF and main ORF start codons on eIF4G2 mRNA. Counts of total 80S ribosome footprints along the 5’ proximal region of the eIF4G2 mRNA after varying durations of 2µg/ml Harringtonine treatment at 37°C. Reads are normalized to library sequencing depth. Curves are smoothened with a 10 nt sliding window.

### A comprehensive view of translation initiation on a single endogenous transcript in vivo

Many of the observations mentioned above derive from metagene analyses which combine data from multiple different transcripts. We asked whether selective 40S footprinting can also be used to gain insights into translation initiation of individual transcripts. To this end, we analyzed eIF4G2, which is interesting because it has both a uORF and a GUG start codon on the main ORF (Fig. 5C). On a 2-dimensional 40S plot, the start codon of the uORF has a characteristic peak with a tail (feature 1, Fig. 5E) indicative of increased 40S dwell-time at this location and start codon recognition. This is accompanied with depletion of eIF2*α*, which is visible on the 1-dimensional 40S plot (feature 2, Fig. 5D), in agreement with some translation initiation on this uORF. Indeed, 80S footprints are visible on the uORF (feature 3, Fig. 5F). Nonetheless, scanning 40S footprints are also visible within the uORF indicating there is also leaky scanning past the uORF start codon (feature 4). On the uORF stop codon 80S footprints decrease and instead eIF2*α* is re-recruited to 40S ribosomes (blue trace, feature 5). All the while, the proportion of 40S ribosomes containing eIF3B, eIF4G1 and eIF4E remains roughly constant, and indeed the scanning 40S ribosomes that can be seen between the uORF and the main ORF are still bound to these initiation factors (feature 6). On the near-cognate GUG start codon of the eIF4G2 main ORF something non-canonical is happening, because the scanning 40S ribosomes recognize the start codon, as can be seen by the high 40S peak at this location (feature 7) and by the characteristic diagonal lines on the 2D plot (feature 8), however eIF2*α* is not depleted (feature 7). This is either because translation is not initiated with eIF2*α* on the GUG start codon, or because start codon recognition is slow due to imperfect pairing with the initiator-tRNA, hence the relative time the 40S resides there prior to eIF2*α* disjoining is increased. This is analyzed more below. Some leaky scanning can be seen by the presence of 40S footprints downstream of the GUG start codon (feature 9), confirming it is a poor initiation signal. Interestingly, these ribosomes become ‘trapped’ by an internal out-of-frame ORF (“iORF”, Fig. 5C), which to our knowledge has not yet been annotated. On this internal ORF initiation starts, as can be seen by a peak on the 40S profile that is depleted of eIF2*α* (feature 10) and the appearance of 80S footprints (feature 11). This gene may have developed these internal out-of-frame ORFs precisely to capture the 40S ribosomes that have scanned past the GUG start codon, to prevent them from scanning down the rest of the mRNA in a cap-tethered fashion, significantly reducing translation efficiency.

We aimed to distinguish between the two possible explanations accounting for the lack of drop in the eIF2*α*-selective 40S footprints on the GUG start codon. The first option is that translation is not initiated on GUG with eIF2*α*. In this scenario, there are two populations of 40S ribosomes – those with eIF2*α* which pause, but scan past the GUG start codon, and those with another unknown initiation factor which initiate translation and convert to 80S ribosomes. In this case, the eIF2*α*-containing 40S ribosomes should become enriched downstream of the start codon, because the other 40S population becomes depleted. To test this, we compared the ratio of eIF2*α*-containing 40S ribosomes to total 40S ribosomes on the GUG peak versus the subsequent peak at position 540nt (feature 9). This shows that roughly the same proportion of ribosomes contain eIF2*α* on the GUG start codon as they do further downstream. This means that if there is another pool of 40S ribosomes containing another initiation factor, it must be a small fraction of the total 40S. To test this, we treated cells with harringtonine, which arrests 80S ribosomes on the start codon. This revealed that the majority of ribosomes initiate on the GUG start codon, and not on the downstream ORFs (Fig 5G). Hence the majority of ribosomes, which are the ones containing eIF2*α*, initiate on the GUG. From this we conclude that translation initiation on this GUG start codon is mediated by eIF2*α*. In sum, selective 40S and 80S footprinting allows a detailed view of the sequential steps of translation initiation occurring *in vivo* and on a single endogenous transcript.

### Co-translational assembly occurs between translation initiation factors of the eIF3 and eIF2 and eIF4F complexes

Up to now we have presented data regarding translation initiation and re-initiation. Selective footprinting also provides insight into events happening during translation elongation. Co-translational assembly is the process whereby a protein interacts with a nascent chain that is still being translated (Shiber et al., 2018). This process is thought to assure the efficient formation of multimeric protein complexes. Co-translational assembly can be observed as an increase in selective footprints when ribosomes are translating the ORF of a gene, due to interaction between the immunoprecipitated protein and the nascent chain encoded by the gene. For example, we observed that eIF3B selective 80S footprints are enriched relative to total 80S footprints on the mRNA of its direct binding partner eIF3A after synthesis of the spectrin domain of eIF3A through which they interact (Fig. 6A) (Dong et al., 2013). Likewise, we find co-translational assembly between eIF3B and eIF3G (Fig. 6B), eIF3B and eIF4G1 (Fig. 6C), eIF2S1 and eIF3A (Fig. 6A), eIF4G1 and eIF3A (Fig. 6A), eIF4E and eIF4G1 (Fig. 6C), and between eIF4E and its export factor eIF4ENIF1 (Fig. 6D). Interestingly, some of these detected interactions are bridged by other factors (eIF4G1 binds eIF3A via eIF3E) and some occur between incomplete protein complexes (eIF4G1 binds eIF3A before it is complete). In summary, we uncover a network of co-translational assembly between the eIF3, eIF4F and eIF2 protein complexes (Figure 6E), suggesting eIF4F may be part of the multi-factor complex in human cells (Sokabe et al., 2012).

**Figure 6:**
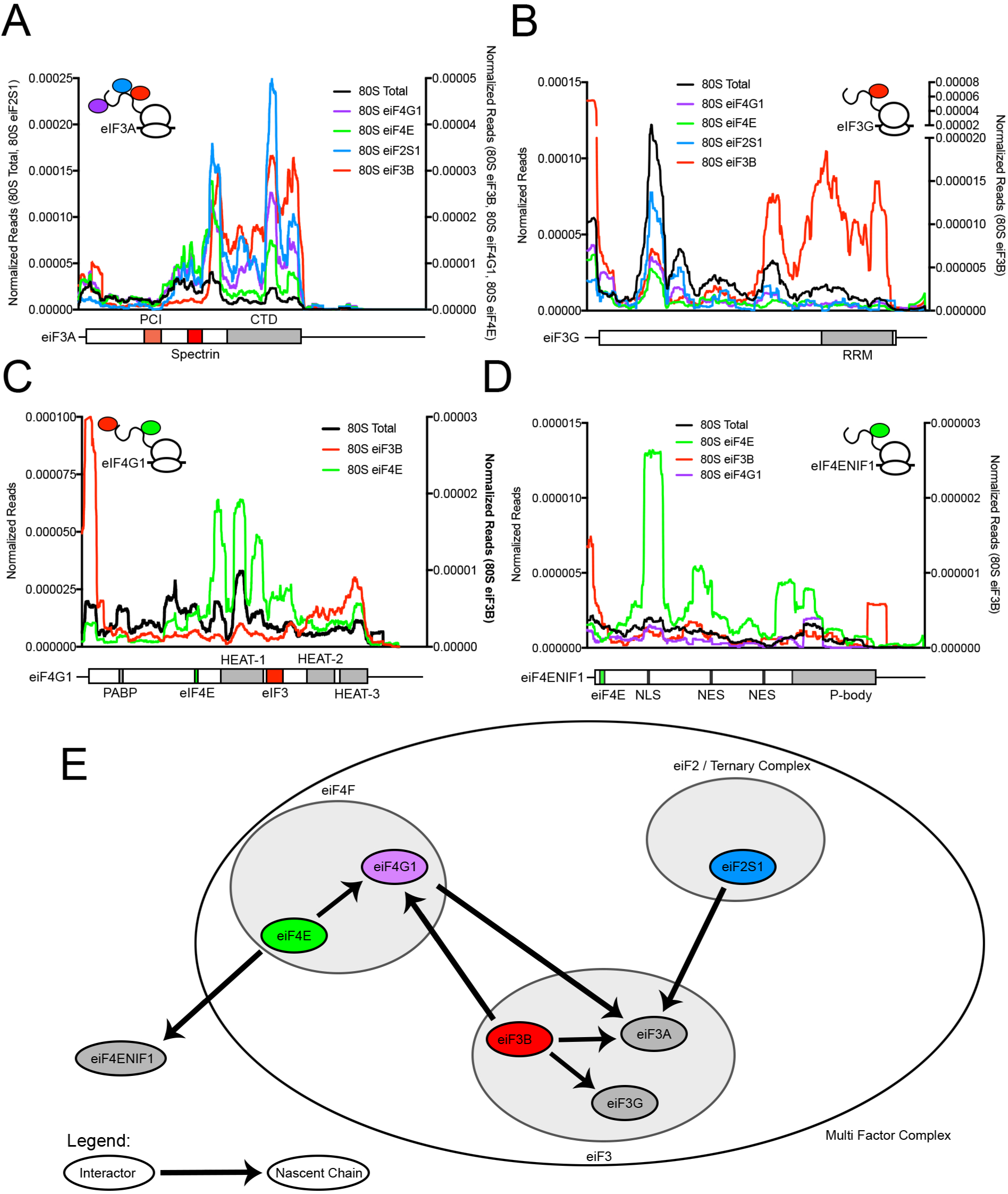
Eukaryotic initiation factor complexes can assemble co-translationally. **(A)** eIF3A assembles co-translationally with eIF2S1, eIF3B, eIF4G1 and possibly eIF4E. Total and selective 80S ribosome footprints on the eIF3A mRNA (ENST00000369144). eIF3B, eiF4E and eIF4G1 selective ribosome footprints are scaled separately. Curves are smoothened with a 200 nt sliding window. The transcript is 6646 nt long. **(B)** eIF3G assembles co-translationally with eIF3B. Total and selective 80S ribosome footprints on the eIF3G mRNA (ENST00000253108). eIF3B selective ribosome footprints are scaled separately. Curves are smoothened with a 50 nt sliding window. The transcript is 1103 nt long. **(C)** eIF4G1 assembles co-translationally with eIF3 and eIF4E. Total and selective 80S ribosome footprints on the eIF4G1 mRNA (ENST00000346169). eIF3B selective ribosome footprints are scaled separately. Curves are smoothened with a 200 nt sliding window. The transcript is 5516 nt long. **(D)** eIF4NIF1 assembles co-translationally with eIF4E. Total and selective 80S ribosome footprints on the eIF4ENIF1 mRNA (ENST00000330125). eIF3B selective ribosome footprints are scaled separately. Curves are smoothened with a 200 nt sliding window. The transcript is 3544 nt long. **(E)** Summary of co-translational assembly of protein complexes and higher order complexes containing eukaryotic initiation factors.

## DISCUSSION

We present here a method that allows the analysis of translation initiation *in vivo* and on endogenous mRNAs in a cell. It allows the investigation of regulation which occurs in a cellular context such as the inactivation of eIF2 in response to ER stress, as we do here. Because the mRNAs are endogenous, they contain the full complement of known and unknown features and modifications that are hard to reproduce in an *in vitro* setting. Hence this will likely serve as a useful approach to complement mechanistic *in vitro* approaches. In the future, it will be interesting to apply this technology to analyzing other aspects of translation initiation. For instance, it will be useful for studying how initiation is altered, and on which mRNAs it is altered, if m6A methylation is reduced, given that the functional role of m6A in translation initiation is a topic of investigation at the moment (Coots et al., 2017; Ozkurede et al., 2019; Patil et al., 2018). Likewise, the function of less-well studied initiation factors such as eIF2A could be revealed.

Using this method, we discover that 1) scanning is mainly cap-tethered in human cells, and 2) that eIF3b, eIF4G1 and eIF4E perdure on translating 80S ribosomes with a half-length of circa 12 codons. This is sterically possible since eIF3 binds the 40S ribosome mainly on the solvent exposed side (Aylett et al., 2015; des Georges et al., 2015) and therefore it does not need to dissociate during 60S subunit joining. These two observations together mean that most ribosomes will remain cap tethered and eIF-associated while scanning and while translating uORFs, which tend to be short, up to the start codon of the main ORF of the mRNA. This has a number of implications. Firstly, it indicates that usually only one ribosome will scan a 5’UTR at a time. This agrees with our harringtonine data where we do not see 40S ribosomes accumulating in front of a stalled initiating 80S ribosome *in vivo* in human cells (Fig. 3D-E and Suppl. Fig. 3G). These data are in contrast to results from rabbit reticulocyte translation extracts where 40S queuing has been observed (Kozak, 1991), suggesting a difference between the *in vitro* and *in vivo* situation. Secondly, this means that 5’ UTR length will influence translation, as we see in our controlled luciferase assays where we modulate only 5’UTR length (Fig. 3F-F’). When comparing endogenous mRNAs with different 5’UTRs lengths in the transcriptome, this trend will be convoluted by the effects of additional sequence elements and secondary structures. Thirdly, it provides a possible explanation for how translation re-initiation in human cells occurs, given that the initiation factors necessary for stabilizing the ribosome’s association with the mRNA and for recruiting a new ternary complex are still present on the ribosome after a uORF. Indeed, *in vitro* studies showed that eIF3 and eIF4F are required for reinitiation with scanning directionality (Poyry et al., 2007; Poyry et al., 2004; Skabkin et al., 2013), and our findings that these initiation factors perdure on early translating 80S ribosomes provides an explanation for how this happens.

The scanning process likely has some quantitative differences in humans compared to yeast. Indeed, ribosomes in yeast are generally not competent to reinitiate after uORFs (Yun et al., 1996). The best studied example where reinitiation happens in yeast is the GCN4 mRNA, where cis-acting elements adjacent to the uORFs are required to stabilize eIF3 on the ribosome and to enable reinitiation (Mohammad et al., 2017; Szamecz et al., 2008). In contrast, mammalian cells do not seem to require cis-acting elements on mRNAs as they are generally reinitiation competent (Johansen et al., 1984; Kozak, 1984; Liu et al., 1984). The difference between the two systems may be quantitative – how strongly ribosomes remain cap-tethered and initiation-factor associated during scanning and elongation. If ribosomes shed initiation factors more quickly in yeast than in humans, for instance after translating 1 or 2 codons, this may render them incapable of uORF reinitiation. An additional difference is that in contrast to what we observe in human cells (Fig. 3F-F’), in yeast 5’UTR length does not affect translation efficiency (Berthelot et al., 2004). Furthermore, 40S queuing *in vivo* has been observed in yeast (Archer et al., 2016). This could be explained if ribosomes in yeast detach from the cap more quickly than in humans, thereby enabling parallel scanning of a 5’UTR by multiple ribosomes. This would likely enable higher translation rates, which would be beneficial for yeast cells that have a rapid cell cycle. Interestingly, the yeast eIF3 complex consists of fewer proteins than the human eIF3 complex (Hinnebusch, 2006; Phan et al., 1998). Furthermore, unlike in humans where the eIF4F complex is linked to the ribosome via eIF3, in yeast it is linked to the ribosome via eIF5 and eIF1 (Asano et al., 2001; He et al., 2003; Shirokikh and Preiss, 2018), which need to leave the ribosome at 60S subunit joining. Thus, it seems likely that eIF4F cannot perdure on translating ribosomes in yeast. Some of these molecular differences may account for quantitative differences in the scanning process.

It is unclear whether *in vivo* mRNAs are circularized, with the 5’cap contacting the 3’ polyA tail via binding of eIF4G to poly-A binding proteins (Adivarahan et al., 2018; An et al., 2018; Gallie, 1991; Khong and Parker, 2018; Wells et al., 1998). It has been postulated that mRNA circularization could help re-recruit terminated ribosomes to the cap for a new round of translation. Our data indicate that 40S ribosomes do not scan down the 3’UTR to the polyA tail after terminating on the main ORF, but rather fall off at the stop codon. Hence, to help re-recruiting ribosomes to the cap, circularization would need to bring the main ORF stop codon close to the cap, either because the 3’UTR is short compared to the main ORF, or because the 3’UTR could be compacted.

In sum, we provide here insights into scanning and translation initiation in human cells. We believe selective 40S footprinting will likely develop into an important powerful approach to study translation initiation mechanisms *in vivo* on endogenous mRNAs in the future.

## MATERIALS & METHODS

### Cloning

Sequences of oligos used for cloning are provided in Supplemental Table 3 at the end of the Materials & Methods. Translation reporters with increasing 5’UTR length were generated as follows. Oligos OJB0440 and OJB0441 were annealed and oligo cloned into a pcDNA3 vector in between a CMV promoter and the Renilla luciferase ORF using HinDIII and Bsp119l sites. The resulting plasmid has a 5’UTR of 60 nt. This plasmid was then opened with EcoR1 and Sal1, separately the 26-mer was excised using EcoR1 and Xho1, and then the 26-mer was inserted into the opened plasmid to double the number of 26-mers in the 5’UTR. This procedure was repeated multiple times to yield the panel of 5’UTR reporters shown in Figure 3 F-F’.

### Cell Culture

HeLa and HEK293T cells were cultured in DMEM +10% fetal bovine serum +100 U/ml Penicillin/Streptomycin (Gibco 15140122). HCT116 cells were cultured in Roti-CELL McCoy’s 5A medium +10% fetal bovine serum +100 U/ml Pencicillin/Streptomycin. Cells were sub-cultured using Trypsin-EDTA for dissociation. Cellular stress conditions were induced by treatment with Tunicamycin at 1 µg / ml or 250 ng / ml. Cells were maintained at 30%-90% confluence.

### Immunoblotting

Protein solutions from sucrose gradients, immunoprecipitations and lysates were run on SDS-PAGE gels and transferred to nitrocellulose membrane with 0.2 µm pore size. After Ponceau staining membranes were incubated in 5% skim milk PBST for 1 hour, briefly rinsed with PBST and then incubated in primary antibody solution (5% BSA PBST or 5% skim milk PBST) overnight at 4°C. Membranes were then washed three times, 15 minutes each in PBST, incubated in secondary antibody solution (1:10000 in 5% skim milk PBST) for 1 hour at room temperature, then washed again three times for 15 minutes. Finally, chemiluminescence was detected using ECL reagents and the Biorad chemidoc. No membranes were stripped.

### Quantitative RT-PCR

Total RNA was isolated from cells using Qiagen RNAeasy spin columns, including on column DNAse digestion to remove plasmid DNA. To synthesize cDNA, 1 µg of total RNA was used for oligo dT primed reverse-transcription using Maxima H Minus Reverse Transcriptase. Quantitative RT-PCR was run with Maxima SYBR Green/ROX mix on a StepOnePlus Real-Time PCR System. Actin B was used as a normalization control and all samples were run in technical triplicates. Non-reverse transcribed RNA, H2O and cDNA from non-transfected Hela cells were assayed to ensure that Renilla luciferase qPCR signal originates from Renilla Luciferase mRNA, not plasmid DNA or unspecific amplification.

### Translation Reporter Dual-Luciferase Assay

For translation reporter assays after plasmid transfection, Hela cells were seeded at 8.000 cells per 96-well. 16-20 hours after seeding, these cells were transfected with three plasmids using lipofectamine 2000. Per well, 60 ng of either GFP expression plasmid, 70 ng of renilla luciferase reporter plasmid and 70 ng of firefly reporter plasmid were used. 3 hours after transfection, the medium was exchanged. Renilla luciferase plasmids always contained the 5’UTR of interest. 0.4 µl Lipofectamine reagent was used per 96-well. Cells were always transfected in six replicates. 16-20 hours after transfection, luciferase activity was assayed using the Dual-Luciferase® Reporter Assay System by Promega according to the manufacturers instructions. To calculate Renilla Luciferase signal per mRNA, only the Rluc signal (not normalized to Fluc) was used and normalized to Rluc mRNA levels from qPCR experiments on cells transfected with the same transfection mixture.

### 40S and 80S Ribosome footprinting

Two days before cell harvest, Hela cells were seeded at 1.5 million cells per 15 cm dish in 20 ml growth medium. For stress conditions, cells were treated with 1 µg/ml or 250 ng/ml tunicamycin 16 hours before cell harvest. For cell harvest, growth medium was poured off and cells were quickly washed with ice-cold washing solution (1x PBS 10 mM MgCl_2_ 800 µM Cycloheximide). Washing solution was immediately poured off and freshly prepared crosslinking solution (1x PBS, 10 mM MgCl_2_, 200 µM Cycloheximide, 0.025% PFA, 0.5 mM DSP) was added to the cells. Cells were incubated with crosslinking solution for 15 minutes at room temperature while slowly rocking. Crosslinking solution was then poured off and remaining crosslinker was inactivated for 5 minutes with ice-cold quenching solution (1x PBS, 10 mM MgCl_2_, 200 µM Cycloheximide, 300 mM Glycine). Quenching solution was poured off and 150 µl of lysis buffer (0,25 M HEPES pH 7.5, 50 mM MgCl2, 1 M KCl, 5% NP40, 1000 μM CHX) was added to each 15 cm dish, resulting in 750µL of lysate. Lysis was carried out at 4°C. Cells were scraped off the dish and lysate was collected. After brief vortexing, lysates were clarified by centrifugation at 20.000g for 10 minutes at 4°C. Supernatant was collected and approximate RNA concentration was determined using a Nanodrop photo-spectrometer. 100 U of Ambion RNAse 1 was added per 120 µg of measured RNA. Lysates were incubated for 5 minutes at 4°C and then loaded onto 17.5%-50% sucrose gradients and centrifuged for 5 hours at 35.000 rpm in a Beckman Ultracentrifuge in the SW40 rotor. Gradients were fractionated using a Biocomp Gradient Profiler system. 40S and 80S fractions were collected for immunoprecipitation and footprint isolation. 40S and 80S fractions corresponding to roughly one or two 15 cm dishes were used for direct extraction of RNA for total footprint samples. 40S and 80S fractions corresponding to roughly ten 15 cm dishes were used for immunoprecipitation of initiation factor bound ribosomes, NP40 was added to these fractions to 1% final concentration. For immunoprecipitation, antibodies were bound to protein A or protein G magnetic dynabeads (Thermo) according to the manufacturers instructions. Antibodies used for ribosome immunoprecipitation are listed in Supplemental Table 5. Beads were washed three times and then added to the 40S or 80S fractions. Fractions with beads were incubated for 2 hours, rotating at 4°C. Then beads were washed three times with bead wash buffer (20 mM Tris pH 7.4, 10 mM MgCl2, 140 mM KCl, 1% NP40), including a change of reaction-vessels during the last wash. Bead volume was increased to ∼500µl with bead wash buffer. Total footprint fractions and IPed fractions were then subjected to crosslink removal and RNA extraction: 55 µl (1/9^th^ of volume) of crosslink-removal solution (10% SDS, 100 mM EDTA, 50 mM DTT) was added, 600 µl Acid-Phenol Chloroform (Ambion) was added and mixture was incubated at 65°C, 1300 rpm shaking for 45 minutes. Tubes were then placed on ice for 5 minutes, spun for 5 min at 20.000 g and supernatant was washed once with acid-phenol chloroform and twice with chloroform, then RNA was precipitated with Isopropanol and subjected to library preparation (see below). The organic phase was used to isolate the precipitated or total proteins. 300 µl Ethanol were added, then 1,5 ml isopropanol were added and solutions were incubated at −20°C for 1 hour. Proteins were sedimented by centrifugation at 20.000g for 20 minutes, washed twice with 95% Ethanol 0,3 M Guanidine HCl, dried and resuspended in 1x Laemmli buffer.

### Deep-sequencing library preparation

During development of the 40S and 80S selective ribosome footprinting method, we optimized and hence changed several parameters. Here we outline first the final, optimized protocol, which we recommend people use for future experiments. Afterwards, we briefly explain the variations of the method which apply to some of the datasets in this manuscript. In Supplemental Table 6 we indicate which deep-sequencing libraries were prepared with which protocol.

Optimized protocol: After RNA extraction from total and IP-purified fractions, RNA quality and integrity were determined on an Agilent Bioanalyzer using the total RNA Nano 6000 Chip. For size selection, RNA was run on 15% Urea-Polyacrylamide gels (Invitrogen) and fragments of size 20-60 nt (80S libraries) and 20-80 nt (40S libraries) were excised using the Agilent small RNA ladder as a reference. RNA was extracted from the gel pieces by smashing the gels into small pieces with gel smasher tubes and extracting the RNA in 0.5 ml of 10 mM Tris pH 7 at 70°C for 10 minutes. Gel pieces were removed and RNA was precipitated using Isopropanol. Footprints were then dephosphorylated using T4 PNK (NEB) for 2 hours at 37°C in PNK buffer without ATP. Footprints were then again precipitated and purified using isopropanol. For 40S footprints, contaminating 18S rRNA fragments were depleted as follows. Prevalent 18S rRNA fragments from the first round of 40S footprinting were used to design complementary Biotin-TEG-DNA oligonucleotides (sequences listed in Supplemental Table 4, ordered from Sigma-Aldrich). 100ng of RNA footprints were then hybridized to a mixture (proportional to occurrence of the fragment, listed in Suppl. Table 4) of these DNA oligos (in 40x molar excess) in (0.5M NaCl, 20mM Tris pH7.0, 1mM EDTA, 0.05% Tween20) by denaturing for 90 sec at 95C and then annealing by reducing the temperature by −0.1C/sec down to 37°C, then incubating 15min at 37°C. Hybridized species were pulled out using Streptavidin magnetic beads (NEB) by incubating at room temperature for 15 minutes, and the remaining RNA was purified by isopropanol precipitation. Footprints were then assayed using an Agilent Bioanalyzer small RNA chip and Qubit smRNA kit. 25 ng or less of footprint RNA was used as input for library preparation with SMARTer smRNA-SeqKit for Illumina from Takara / Clontech Laboratories according to the manufacturer’s instructions. Deep-sequencing libraries were sequenced on the Illumina Next-Seq 550 system.

For RNA-seq libraries, total cell RNA was extracted using TRIzol and library preparation was performed using the Illumina TruSeq Stranded library preparation kit. These RNA-seq libraries were also sequenced on the Illumina Next-Seq 550 system.

Protocol variants used for some of the datasets: Two 80S footprint libraries and one RNA-seq library (as listed in Suppl. Table 6) were prepared using a the NEXTflex Small RNA-Seq Kit v3 library preparation kit instead of the Takara/Clonentech kit. For this, size selected rRNA depleted RNA was phosphorylated using T4 PNK. Footprints were then assayed using an Agilent Bioanalyzer small RNA chip and a Qubit smRNA kit. Deep sequencing libraries were prepared from these RNA fragments using the Bio-Scientific NEXTflex Small RNA-Seq Kit v3. Deep-sequencing libraries were sequenced on the Illumina Next-Seq 550 system.

In some cases (Suppl. Table 6), the Ribo Zero Gold rRNA depletion kit (Illumina) was used instead of our custom rRNA depletion protocol mentioned above. However, this kit was discontinued half-way though the project and is no longer available.

### Data Analysis

Adapter sequences and randomized nucleotides (Nextflex) or polyA stretches (Clonentech) were trimmed from raw reads using cutadapt. Nuclear and mitochondrial Ribosomal RNA and tRNA reads were removed by alignment to human tRNA and rRNA sequences using bowtie2. Then, the remaining reads were separately aligned to the human transcriptome (Ensemble transcript assembly 94) and human genome using BBmap. Multiple mappings were allowed. Secondary mappings were counted when analyzing single transcripts, but not counted in metagene plots to avoid biasing genes with many transcript isoforms or reads with low sequence complexity. Metagene plots, single transcript traces and grouped analyses were carried out or created with custom software written in C, supplied in Supplemental Dataset 1. Read counts for metagene plots of whole transcripts that encompass 5’UTRs, ORFs and 3’UTRs (Fig. 1B) were normalized for the length of each of these features to make them comparable. 70 of the 41.314 transcripts were excluded from the analyses (listed in Suppl. Table 8) because PCR artefacts mapped to these transcripts. 2-D metagene plots were visualized using Fiji (Schindelin et al., 2012). For 80S plots, only reads with footprint lengths between 26 to 37 nt were counted.

Translated uORFs were defined by the presence of any 80S ribosome footprints in a 10-nucleotide window around the uORF start codon. Only ATG initiated uORFs were considered.

### Data and software availability

All custom software used in this study is supplied as Supplemental Dataset 1. All library sequencing data are available at NCBI Geo (we are in the process of submitting it, and the accession number will be available at revision stage). A table summarizing read counts per transcript for different experiments is provided as Supplemental Table 7.

## ACKNOWLEDGEMENTS

We thank Georg Stoecklin, Johanna Schott and Kathrin Leppek for sharing with us their ribosome footprinting protocol, for scientific discussion, and for giving us the eIF3B antibody. We thank Oliver Hoppe for designing the graphical abstract. This work was funded in part by a Deutsche Forschungsgemainschaft (DFG) Collaborative Research Center SFB1036 grant to B.B. and A.A.T., by a DKFZ NCT3.0 Integrative Project in Cancer Research (NCT3.0_2015.54 DysregPT) grant to B.B. and A.A.T., and by a Cell Networks – Cluster of Excellence (EXC81) grant to J.B. High-throughput sequencing was carried out at the DKFZ Genomics and Proteomics Core Facility, or using an Illumina NextSeq 550 system funded by the Klaus Tschira Foundation gGmbH, Heidelberg, Germany.

## AUTHOR CONTRIBUTIONS

Experiments were performed by J.B. All authors designed the work, analyzed data, interpreted data, and wrote the manuscript.

---

**Supplemental Figure 1:**
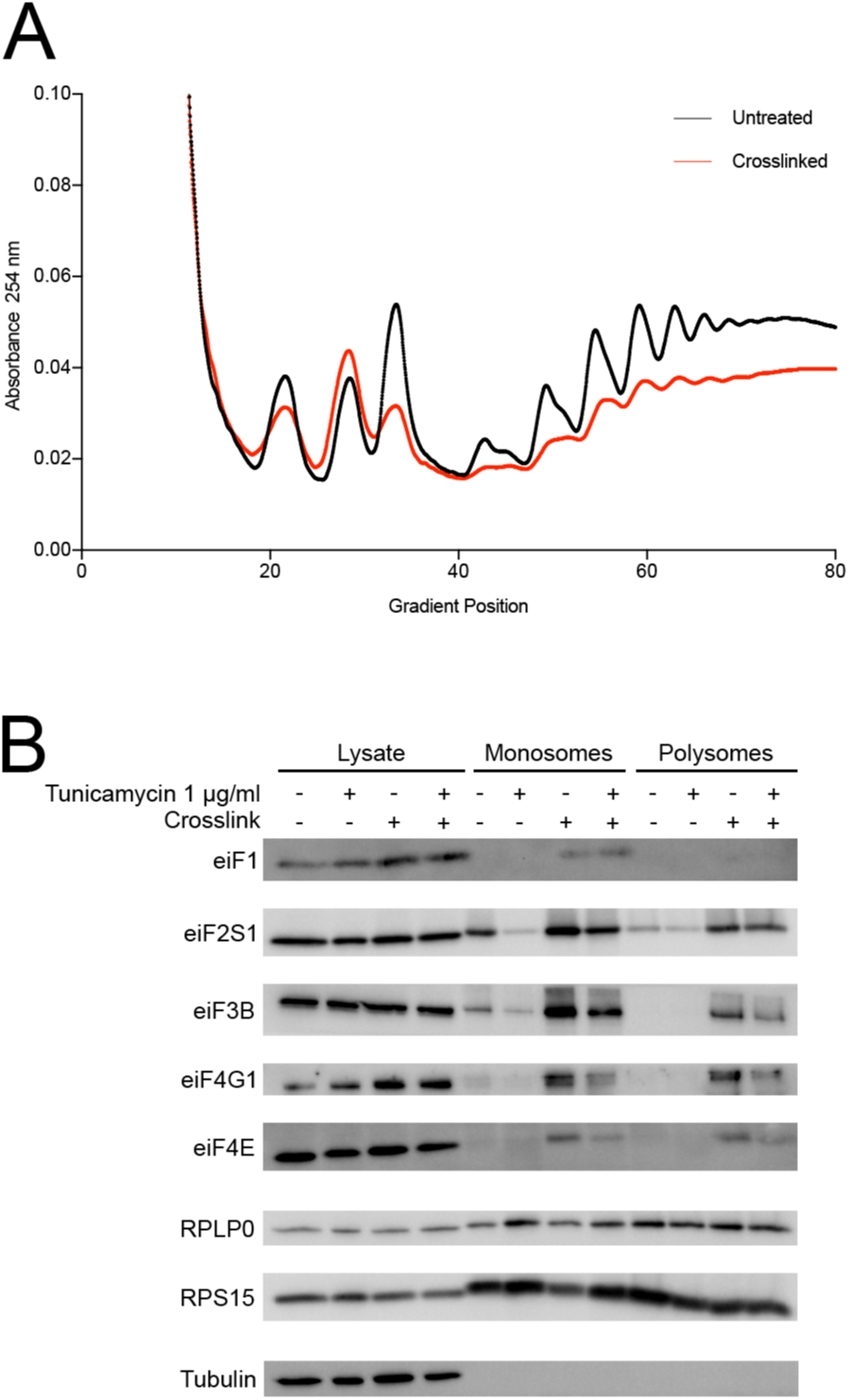
Crosslinking with 0.025% PFA + 0.5 mM DSP strikes a compromise between sufficient retention of initiation factors and 40S ribosomes on mRNAs while avoiding non-specific aggregate formation. **(A)** Crosslink protocol causes mild decrease in polysome yield. Polysome gradients of HeLa cells treated with or without 0.5 mM DSP+0.025% para-formaldehyde. **(B)** Crosslink stabilizes interaction of eukaryotic initiation factors with ribosomes. Western blot of sucrose gradient fractions from (A) pooled into monosomes and polysomes. Additionally, cells were treated with 1 µg/ml tunicamycin or left untreated.

**Supplemental Figure 2:**
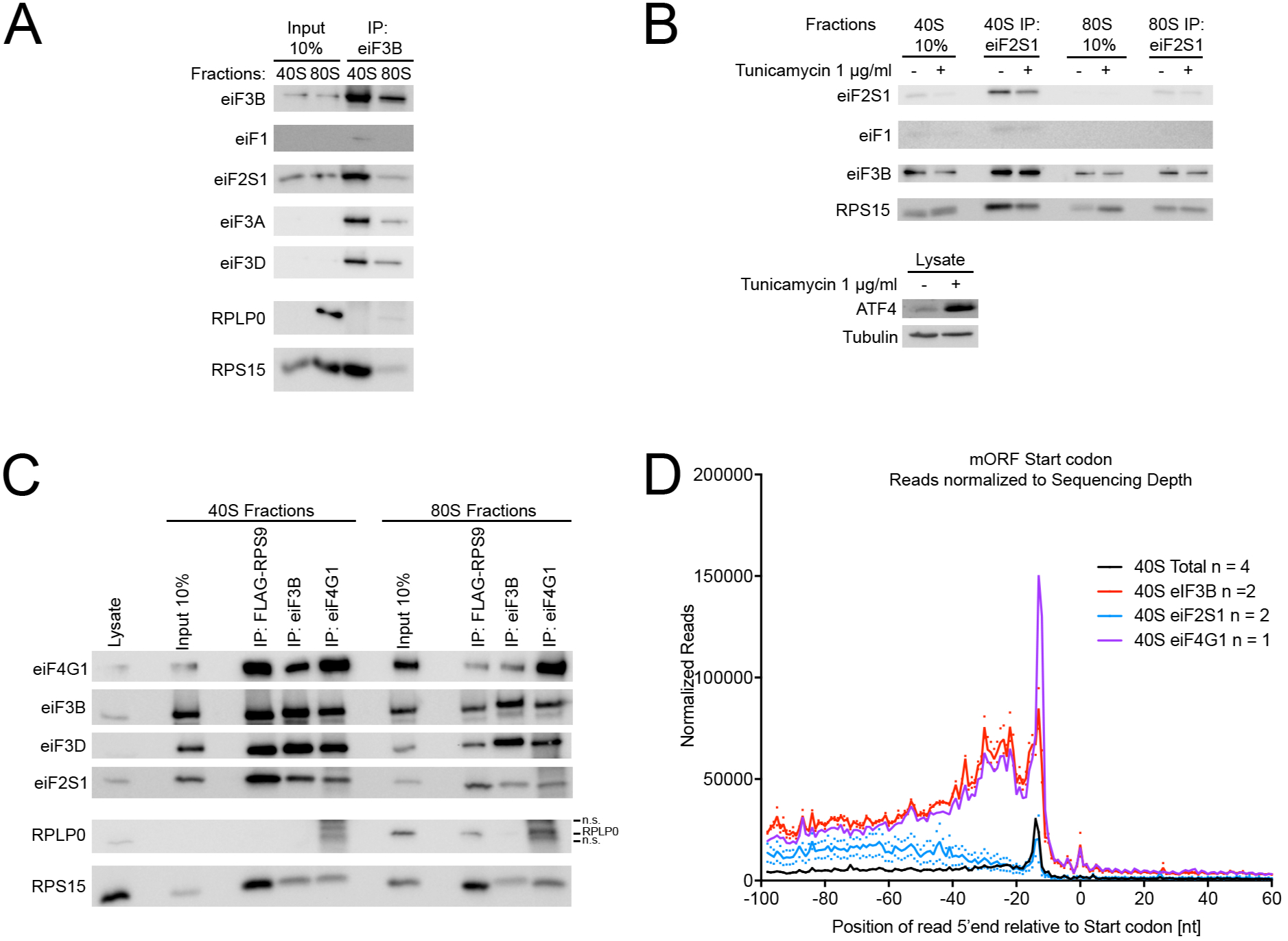
Selective ribosome footprinting for eIF2S1 and eIF3B associated ribosomes. **(A)** Immunoprecipitation of eIF3B from 40S and 80S ribosomes. Preparative IP of eIF3B from 40S and 80S fractions after crosslinking, RNAse digestion and sucrose gradient centrifugation. **(B)** Immunoprecipitation of eIF2S1 from 40S and 80S ribosomes. Preparative IP of eIF2S1 from 40S and 80S fractions after crosslinking, RNAse digestion and sucrose gradient centrifugation. Cells were either untreated of stressed with 1 µg/ml tunicamycin for 16 hours. **(C)** Immunoprecipitation of eIF3B and eIF4G1 from 40S and 80S ribosomes. Preparative IP of eIF3B and eIF4G1 from 40S and 80S fractions after crosslinking, RNAse digestion and sucrose gradient centrifugation. FLAG-RPS9 precipitation was not used for sequencing because of a high background of non-specific ribosome binding in the purification. **(D)** Start codon metagene plot of total and eIF3B and eIF2S1 selective 40S ribosome footprints on all human protein coding transcripts (n = 41.244). Reads are mapped to the position of the read 5’ end. Graphs are normalized to the number of mapped reads in each library. These are the same data as in main Figure 2A, but normalized only for library size.

**Supplemental Figure 3:**
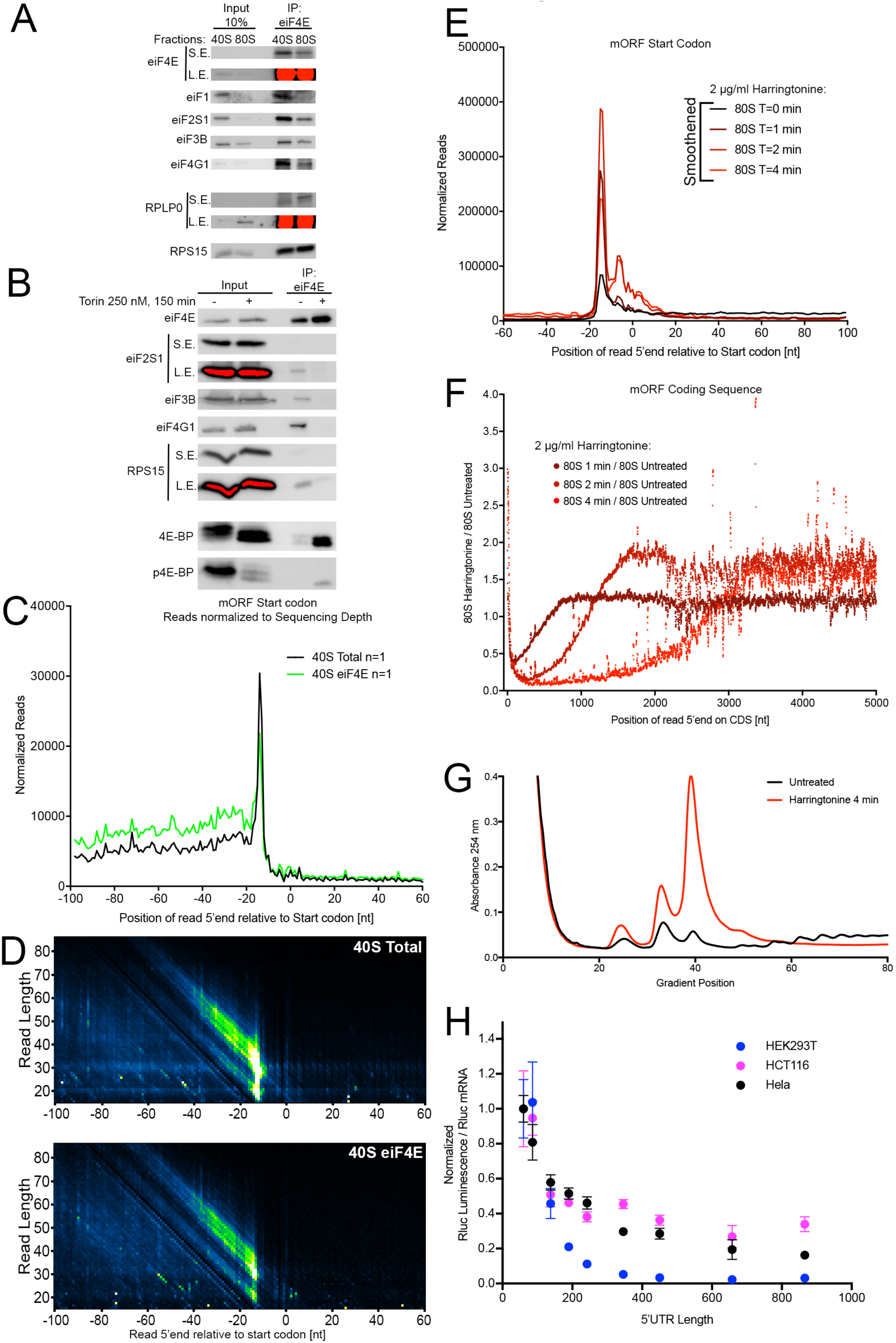
Selective ribosome footprinting for eIF4G1 and eIF4E associated ribosomes. **(A)** Immunoprecipitation of eIF4E from 40S and 80S ribosomes. Preparative IP of eIF4E from 40S and 80S fractions after crosslinking, RNAse digestion and sucrose gradient centrifugation. L.E.=long exposure, S.E.=short exposure **(B)** Co-IP of ribosomes with eIF4E is specific, as it significantly decreases upon mTOR inhibition. HeLa cells were treated with 250 nM Torin for 150 minutes or with vehicle, then crosslinked, lysed and RNAse treated, then eIF4E was immunoprecipitated. L.E.=long exposure, S.E.=short exposure **(C-D)** Start codon metagene plot of total and eIF4E-selective 40S ribosome footprints for all human protein coding transcripts (n = 41.244). Reads are mapped to the position of the read 5’ end. Graphs are normalized to the number of mapped reads in each library. 2-dimensional plot resolving for footprint length shown in (D). **(E-F)** Harringtonine treatment was successful because it caused accumulation of 80S ribosomes on start codons and run-off of translating 80S ribosomes from Open Reading Frames. (E) Start codon metagene plot of total 80S ribosome footprints after varying times of 2 µg/ml Harringtonine treatment for all human protein coding transcripts (n = 41.244). Reads are mapped to the position of the read 5’ end. Curves are normalized to the total number of mapped reads in each library. (F) Metagene plot showing the ratio of 80S ribosome footprints in harringtonine-treated cells versus untreated control cells at various positions along the coding sequence of all protein coding mRNAs (n = 41.244). Read 5’ end is mapped and offset by +15 nucleotides. Read counts are normalized to number of mapped reads in each library. Curves were smoothened with sliding window = 10 nt. **(G)** Sucrose density gradient of cell lysates of crosslinked cells that were either treated with 2 µg/ml Harringtonine for 4 minutes or untreated. **(H)** 5’UTR-length limits the translational output of mRNAs in various human cell lines. Renilla luciferase luminescence normalized to mRNA levels for translation reporters containing 5’UTRs of varying lengths shown in main Figure 3F. Renilla luciferase mRNA levels were normalized to Actin B mRNA levels. Representative experiments of at least two biological replicates are shown.

**Supplemental Figure 4:**
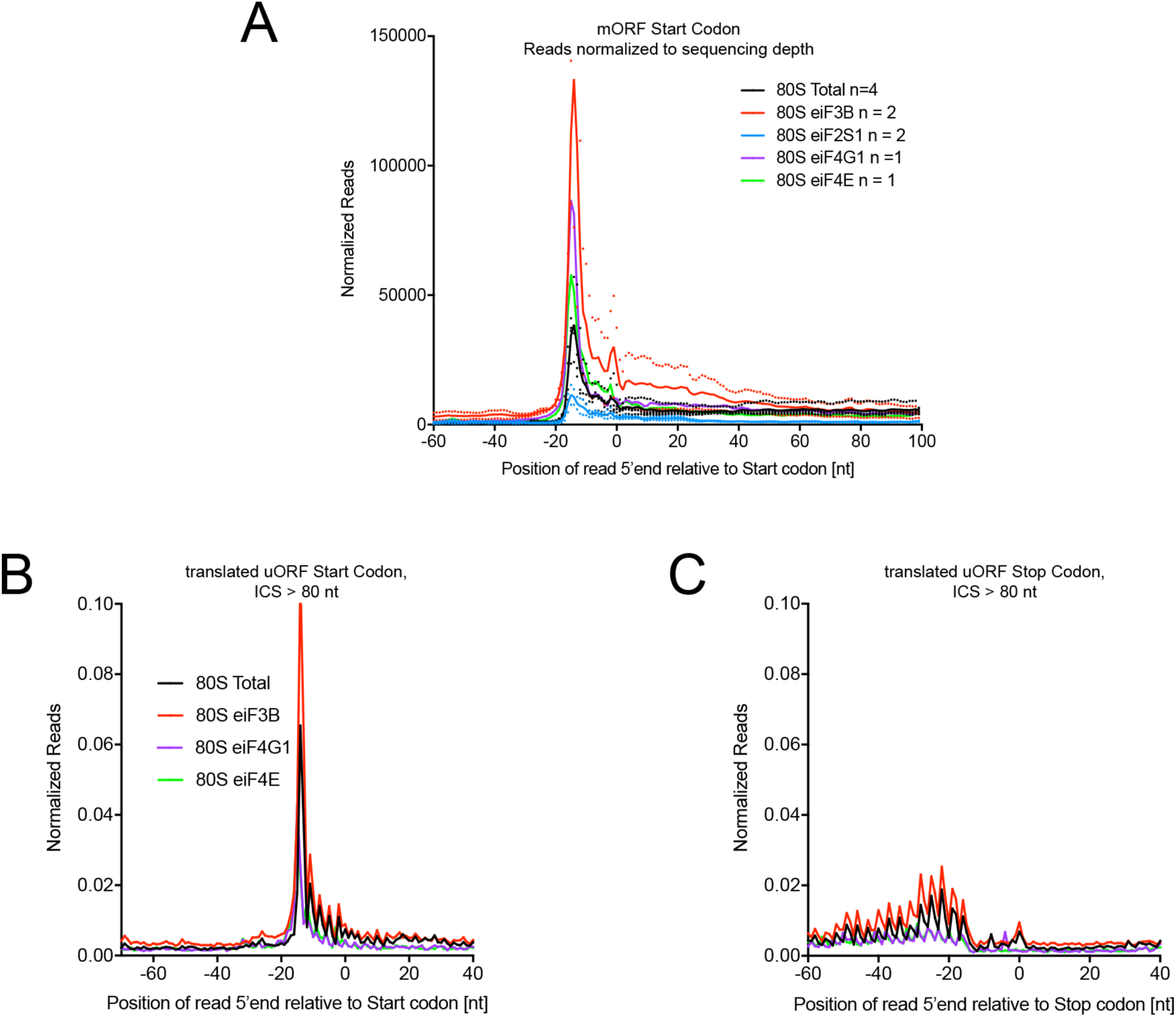
Selective 80S ribosome footprinting for eIF3B eiF2S1, eIF4G1 and eIF4E associated ribosomes. **(A)** Start codon metagene plot of total and selective 80S ribosome footprints on all human protein coding transcripts (n = 41.244). Reads are mapped to the position of the read 5’ end. Same data as in main Figure 4A, except here the curves are only normalized to the number of mapped reads in each library. **(B-C)** 80S ribosomes inside translated uORFs exhibit triplet periodicity. Metagene plots of total and selective 80S footprints relative to the start codon (B) or stop codon (C) of translated uORFs (n = 6663). Reads are mapped to the position of the read 5’ end. Curves are normalized to each other using the scaling factors from Figure 4A (for 80S). Single replicates are shown here.

