## Supplementary Information for "Selective 40S footprinting reveals that scanning ribosomes remain cap-tethered in human cells"

Bohlen et al.

### Contents

1) Supplemental Tables ........................................................................... p. 1

2) List of custom software provided in Suppl. Dataset 1 ...................................... p. 4

### Supplemental Tables

Supplemental Table 1: Antibodies used for immunoblotting

| Antigen | Provider and Product Number | Blotting Dilution |
|---|---|---|
| Tubulin | T9026 Sigma | 1:2500 |
| RPS15 | Atlas HPA054510 | 1:250 |
| RPLP0 | Atlas HPA003512 | 1:250 |
| eiF2S1 | Cell Signaling #5324 | 1:1000 |
| eIF3B | Santa Cruz 16377 (Discontinued) | 1:1000 |
| eIF3A | Cell Signaling #3411 | 1:1000 |
| eIF4E | MBL RN001P | 1:1000 |
| eIF4G1 | MBL RN002P | 1:1000 |
| eIF1 | OriGene TA502844S | 1:1000 |
| eIF3D | Atlas HPA063330 | 1:1000 |

| | | |
|---|---|---|
| ATF4 | Cell Signaling #11815 | 1:500 |
| anti phospho 4E-BP1 (Ser65) | Cell Signaling #9451 | 1:1000 |
| 4E-BP1 | Cell signaling 9452 | 1:1000 |

Supplemental Table 2: q-RT-PCR Primers

| **Target** | **Sequence** |
|---|---|
| Actin B forward | CCACCATGTACCCTGGCATT |
| Actin B reverse | CGCTCAGGAGGAGCAATGAT |

Supplemental Table 3: Cloning Oligos

| **Name** | **Sequence** | **Purpose** |
|---|---|---|
| OJB0440 | AGCTTgaattcgatcgtcgacATCTCACGACCCAC TACACACTCGAGCCACCATGACTT | 5'UTR length oligo cloning Forward (HindIII BSP119L) |
| OJB0441 | CGAAGTCATGGTGGCTCGAGTGTGTAGTG GGTCGTGAGATgtcgacgatcgaattcA | 5'UTR length oligo cloning Reverse (HindIII BSP119L) |

Supplemental Table 4: 18S rRNA depletion Oligos

| **Oligo#** | **Sequence** | **Proportion** |
|---|---|---|
| 1 | GGGGGTCAGCGCCCGTCGGCATGTATTAGCTCTAG AATTACCACAGTTATCCAAGTA | 21x |
| 2 | GTGCGATCGGCCCGAGGTTATCTAGAGTCACCAAA GCCGCCGGCGCCCGCCCCCCGGCCG | 16x |
| 3 | TTACGACTTTTACTTCCTCTAGATAGTCAAGTTCGAC CGTCTTCTCAGCG | 1x |
| 4 | AGCGAGCGACCAAAGGAACCATAACTGATTTAATGA GCCATTCGCAGTTTCACTGTAC | 2x |
| 5 | GGGCGCCGAGAGGCAAGGGGCGGGGACGGGCGG TG | 2x |
| 6 | CGTACTTAGACATGCATGGCTTAATCTTTGAGACAA GCATATGCTACTGGCAGGATCAACCAGGTA | 1x |
| 7 | GCCGGAGAGGGGCTGACCGGGTTGGTTTTGATCTG ATAAATGCACGCATCCCCCCCG | 1x |
| 8 | CGACTACCATCGAAAGTTGATAGGGCAGACGTTCG AATGGGTCGTCGCCG | 1x |

| | | |
|---|---|---|
| 9 | GTTCGTCCAAGTGCACTTTCCAGTACACTTACCATG TTACGACTTGTCTCCTCT | 1x |
| 10 | AGCCTCGCCCTGGGAAAACCACCTTCGTGATCATG GTATCTCCCCTGCCA | 1x |
| 11 | GGCCTGCTTTGAACACTCTAATTTTTTCAAAGTAAAC GCTTCGGGCCCCG | 1x |
| 12 | GAAAACATTCTTGGCAAATGCTTTCGCTCTGGTCCG TCTTGCGCCGGTCC | 1x |
| 13 | GCTGCGTTCTTCATCGACGCACGAGCCGAGTGATC CACCGCTAAGAGTCG | 1x |
| 14 | ATTCCATTATTCCTAGCTGCGGTATCCAGGCGGC | 1x |
| 15 | ACCCCGGCGGGGCCGATCCGAGGGCCTCACTAAA CCATCCA | 1x |
| 16 | TTGCCCTCCAATGGATCCTCGTTAAAGGATTTAAAG TGGACTCATTCCAATTACA | 1x |
| 17 | CGCCGGCCGAGGTGGGATCCCGAGGCCTCTCCAG TCCGCCGAGGGCGCACCACCGGCCCGTCTCGC | 1x |
| 18 | TCTTAATCATGGCCTCAGTTCCGAAAACCAACAAAA TAGAACCGCGGTCC | 1x |
| 19 | CTCCCCGGGTCGGGAGTGGGTAATTTGCGCGCCTG CTGCCTTCCTTGGAT | 1x |
| 20 | AATAACGCCGCCGCATCGCCGGTCGGCATCGTTTA TGGTCGGAACTACGACGGTATC | 1x |

Supplemental Table 5: Antibodies used for ribosome immunoprecipitation

| **Antigen** | **Provider and Product Number** | **Used amount for one sequencing library** |
|---|---|---|
| eIF3B | Santa Cruz 16377 (Discontinued) | 40 µl |
| eIF2S1 | Proteintech 11170-1-AP | 70 µl |
| eIF4G1 | MBL RN002P | 40 µl |
| eIF4E | MBL RN001P | 75 µl |

Supplemental Table 6: List of libraries generated and sequenced in this study.

(provided as a separate Excel file)

Supplemental Table 7: Read counts per transcript for the experiments presented in this paper.

(provided as a separate Excel file)

Supplemental Table 8: List of transcripts excluded from the analyses

(provided as a separate Excel file)

### List of custom software provided in Suppl. Dataset 1

**count_features_nt_range_v2.c**

*Description*: Takes as input a SAM file with mapped reads, and a file containing a list of transcripts including a nucleotide range for each transcript, and the software counts the number of reads that fall within the nucleotide range of each transcript. Secondary mappings are excluded.

*Inputs*:

•the name of the SAM file to be processed should be indicated as a command line argument

•a tab-delimited file called "in_features.txt". No header row. First column: transcript name (to match the transcript name in the SAM file). Second column: nucleotide range start. Third column: nucleotide range stop.

*Output*: a tab-delimited file with read counts per transcript. Progress of the program is shown to the user by displaying a dot (‘.’) on the screen every 10000 lines processed.

**metagene_plot_5UTR_CDS_3UTR_v2.c**

*Description:* Performs the counting to generate a combined metagene plot of reads in 5’UTR, ORF and 3’UTR of transcripts. Read counts were normalized for the length of each of these features to make them comparable (yielding a measurement of ‘read density’).

*Inputs*:

•the name of the SAM file to be processed should be indicated as a command line argument

•a tab-delimited file called “in_features.txt”. No header row. First column: transcript name (to match the transcript name in the SAM file). Second column: total length of the transcript in nucleotides. Third column: position of the first nucleotide of the start codon of the main Open Reading Frame on the transcript. Fourth column: position of the last nucleotide of the stop codon of the main Open Reading Frame on the transcript.

*Output*: a tab-delimited file with read counts per position, first for 5’UTRs, then ORFs, then 3’UTRs. Progress of the program is shown to the user by displaying a dot (‘.’) on the screen every 10000 lines processed.

**metagene_plot_interval_v6_footprint_len.c**

*Description:* Performs the counting to generate a 2-dimensional metagene plot of reads within an interval, with position in that interval normalized to 0-100% (e.g. all transcript 5'UTRs), resolved for footprint length (1nt-100nt). Secondary mappings are not counted to avoid biasing genes with many transcript isoforms or reads with low sequence complexity. Only one interval per transcript is allowed.

*Inputs*:

•SAM file entitled “input.sam” to be processed

•a tab-delimited file called “in_features.txt”. No header row. First column: transcript name (to match the transcript name in the SAM file). Second column: interval start position in nucleotides. Third column: interval end position in nucleotides.

*Output*: a tab-delimited file ‘output.txt’ with read counts per position percentile (in rows), resolved for footprint length (in columns). Progress of the program is shown to the user by displaying a dot (‘.’) on the screen every 10000 lines processed.

**metagene_plot_onePoint_v7_footprint_len.c**

*Description:* Performs the counting to generate a 2-dimensional metagene plot of reads relative to fixed point (e.g. start codon), from 100nt upstream of the fixed point to 100nt downstream, resolved for footprint length (1nt-100nt). Secondary mappings are not counted to avoid biasing genes with many transcript isoforms or reads with low sequence complexity. Only one interval per transcript is allowed.

*Inputs*:

•SAM file entitled “input.sam” to be processed

•a tab-delimited file called “in_features.txt”. No header row. First column: transcript name (to match the transcript name in the SAM file). Second column: position of the fixed point, in nucleotides, for that transcript.

*Output*: a tab-delimited file ‘output.txt’ with read counts per position (in rows), resolved for footprint length (in columns). Progress of the program is shown to the user by displaying a dot (‘.’) on the screen every 10000 lines processed.

**single_gene_plot_v2.c**

*Description:* Counts the number of reads as a function of position on a single transcript, resolved for footprint length (1nt-100nt). Secondary mappings are counted.

*Inputs*:

•the name of the SAM file to be processed should be indicated as the first command line argument

•the name of the transcript to be analysed should be indicated as the second command line argument

•a tab-delimited file called “in_features.txt” containing information for all transcripts, as for the program “metagene_plot_5UTR_CDS_3UTR_v2.c”. No header row. First column: transcript name (to match the transcript name in the SAM file). Second column: total length of the transcript in nucleotides. Third column: position of the first nucleotide of the start codon of the main Open Reading Frame on the transcript.

Fourth column: position of the last nucleotide of the stop codon of the main Open Reading Frame on the transcript.

*Output*: a tab-delimited file ‘output.txt’ with read counts per position (in rows), resolved for footprint length (in columns). Progress of the program is shown to the user by displaying a dot (‘.’) on the screen every 10000 lines processed.
